# Ocrelizumab Modulates Both B and T Cell Immune Capacities in Multiple Sclerosis

**DOI:** 10.64898/2026.03.24.713043

**Authors:** Qi Wu, Mikel Gurrea-Rubio, Qin Wang, Deanna Dwyer, Elizabeth A Mills, Josh Garton, Joshua S Mytych, Steven K Lundy, Christopher D Scharer, Jeremy Boss, Laura Cooney, Dean E Draayer, Phillip L Campbell, David A. Fox, Yang Mao-Draayer

## Abstract

To understand the molecular and cellular mechanisms beyond B-cell depletion with the anti-CD20 monoclonal antibody ocrelizumab, we used comprehensive muti-modal flow cytometry and functional assays in a prospective longitudinal multiple sclerosis (MS) cohort. Ocrelizumab depleted the vast majority of B cells and showed selective effects on different B cells subsets. Analysis of residual/replenished B cells revealed relative enrichment of regulatory B cells like CD27^+^CD43^+^ B1 and CD24^hi^CD38^hi^ transitional B cells, and reduction of CD27^+^ memory B cell subsets and CD19^+^IgD^+^CD27^−^naïve B cells at early time points (1-3 month) and before subsequent infusions at 4-7 months, 11-14 months, and >18 months. CD20+ T cells and peripheral helper T-cells (Tph) were also reduced. RNA sequencing analysis showed B1 cells have significantly higher expression of *LGALS1*, *KCNN4*, *ITGB1*, and *IL2RB*. Compared to transitional B cells, B1 cells also displayed significantly higher expression of tissue homing molecules *ITGAX (CD11c)*, *S100A4*, *ITGB1*, and *CXCR3*. IL10 signaling pathway is increased in these B cells. Ex vivo B cell functional assays indicated the residual/replenishing B cells were anergic following ocrelizumab, with increased IL10/TNFα and IL10/IL6 ratios under BCR stimulation. Ocrelizumab treatment may create a self-reinforcing regulatory circuit: the reduction of Tph cells could alleviate suppression of regulatory B cells, which subsequently expand and further promote regulatory T cell networks via *IL2RB*, *LGALS1*, and an increased IL-10 signaling pathway.

## Introduction

Multiple sclerosis (MS) is the most common demyelinating disease of the central nervous system (CNS), mainly affecting young adults^1, 2^. MS is characterized by episodes of relapse with complete or incomplete remissions, eventually leading to permanent neurological impairment in many patients. T and B lymphocytes are critical cellular mediators in the pathophysiology of MS, but disease activity is differentially impacted across clinical subtypes, with some subsets of lymphocytes driving damage while other subsets facilitate repair. However, our current understanding of the most relevant pathogenic and protective subsets is incomplete. Effective disease-modifying treatments (DMTs) must both limit unwanted immune responses associated with disease initiation and propagation, as well as have minimal adverse impact on normal protective immune responses^3^. Therefore, understanding the mechanism of action (MOA) of efficacious DMTs may provide insight into the underlying pathophysiology of MS.

Ocrelizumab is a humanized monoclonal antibody that targets CD20 on pre-B, immature and mature B cells^4^. Ocrelizumab was shown to have therapeutic benefits for progressive MS patients, yet the mechanism by which it confers this benefit is not known. In experimental autoimmune encephalomyelitis (EAE), a mouse model of MS, it was shown that pathogenic B cells persist after anti-CD20 antibody treatment^5^; however, less is known about the corresponding T and B cell changes in MS patients. CD20 is not expressed by antibody-secreting plasmablasts and plasma cells^6^, this suggests that the pathogenic role of B cells in MS is not confined to the generation of autoantibodies but also includes their capacity to produce cytokines and to interact with other immune cell types, such as T cells^7^.

Aside from pathogenic B cells, several subsets of specialized IL10-producing B cells (Bregs) have been identified in humans, including transitional and B1 cells^8–13^. While these subsets may harbor regulatory potential and play a role in mediating tolerance^11^, the populations differ greatly in terms of localization, differentiation, and proposed function. B1 (CD27^+^CD43^+^) cells are innate-like cells, derived largely from fetal and neonatal B cell lymphopoiesis by progenitors distinct from those of conventional B cells^14^, ^14^, and are associated with natural IgM production^16^. CD27^+^CD24^hi^ B cells comprise another potential IL-10 producing subset, which has been proposed as equivalent of murine B10 cells. They produce IL-10, TGFβ, and TNFα and can suppress CD4 T cell proliferation and production of IFNγ and IL-17. Transitional B cells (TransB) are characterized by the cell surface markers CD24^hi^CD38^hi^ in humans and represent an immature B cell population found in the peripheral blood that produce IL-10^17^. They influence the balance of T cells toward a more anti-inflammatory phenotype through the suppression of Th1 and Th17 cell differentiation, enhancement of Th2 polarization, and induction of regulatory T cells (Tregs)^18, 18^. These B cell subsets are associated with immune regulation in the context of autoimmune diseases.

To understand the impact of ocrelizumab treatment on these and other immune cell subsets, we performed extensive immunophenotyping and functional assays before and throughout treatment to include not only short-term effects at 1-3 months and 4-7 months, but also long-term effects at 11-14 months and >18 months. We found that ocrelizumab not only significantly reduced the B cell populations, but also CD20^+^ T cells, and especially peripheral helper T-cells (Tph). We showed that ocrelizumab has differential effects on B cell subsets - while the frequency of naïve B and unswitched memory B cells is decreased, the frequency of transitional and B1 cells is significantly increased with ocrelizumab. We further sorted various B cell subsets for transcriptomic analyses in order to test the validity of assumptions concerning their functional programs. In addition, we performed ex vivo functional assays on ocrelizumab-treated MS patients’ B-cells and showed that they exhibit an exhausted phenotype and an increased anti-inflammatory capacity with an increased ratio of IL-10/IL-6 and IL-10/TNFα.

## Material and Methods

### Participant demographics

All enrolled patients were from the Autoimmunity Center of Excellence and Multiple Sclerosis Center at the University of Michigan. Recruitment and initial eligibility screening were accomplished by medical record review. Prior to participation in the study, informed consent, which was approved by the University of Michigan Institutional Review Board, was obtained from each patient. The study was conducted according to the Declaration of Helsinki and in accordance with good clinical practice guidelines. The inclusion criteria were all subjects (age 18-65) with RRMS. Refer to clinicaltrials.gov identifier NCT04459988 for more details. Blood samples were collected at baseline (pre-treatment) and longitudinally followed post treatment at 1-3 months, as well as at 4-7 months, 11-14 month and >18 months prior to the following infusions. The demographic characteristics of participants in the study are summarized in Table 1. The cohort included both naïve patients and patients who switched from other DMTs.

**Table 1.**
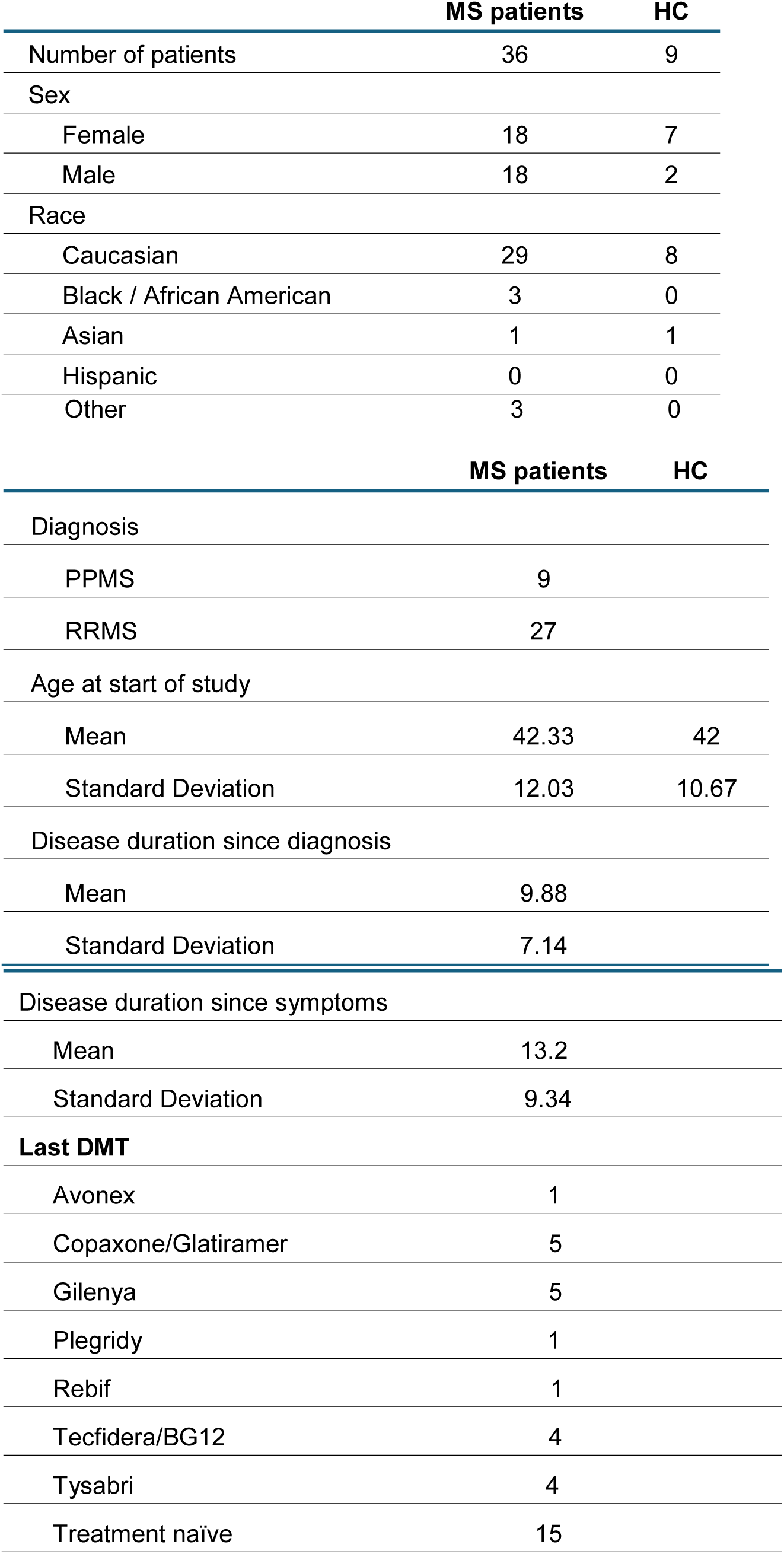
Demographics of MS and health (HC) subjects recruited for this study.

|  | MS patients | HC |
| --- | --- | --- |
| Number of patients | 36 | 9 |
| Sex |  |  |
| Female | 18 | 7 |
| Male | 18 | 2 |
| Race |  |  |
| Caucasian | 29 | 8 |
| Black / African American | 3 | 0 |
| Asian | 1 | 1 |
| Hispanic | 0 | 0 |
| Other | 3 | 0 |
|  | MS patients | HC |
| Diagnosis |  |  |
| PPMS | 9 |  |
| RRMS | 27 |  |
| Age at start of study |  |  |
| Mean | 42.33 | 42 |
| Standard Deviation | 12.03 | 10.67 |
| Disease duration since diagnosis |  |  |
| Mean | 9.88 |  |
| Standard Deviation | 7.14 |  |
| Disease duration since symptoms |  |  |
| Mean | 13.2 |  |
| Standard Deviation | 9.34 |  |
| **Last DMT** |  |  |
| Avonex | 1 |  |
| Copaxone/Glatiramer | 5 |  |
| Gilenya | 5 |  |
| Plegridy | 1 |  |
| Rebif | 1 |  |
| Tecfidera/BG12 | 4 |  |
| Tysabri | 4 |  |
| Treatment naïve | 15 |  |

Patient data were obtained through medical records at the University Hospital, Michigan Medicine, Ann Arbor, MI. Routine CBC with differential blood labs were done at UM hospital clinical lab. Other immune assays (described below) were done in the UM NIH NIAID funded Autoimmunity Center of Excellence (ACE) lab.

### Peripheral blood mononuclear cell (PBMC) isolation

High-quality PBMCs were isolated from all healthy control (HC) participants and from MS patients prior to and following treatment initiation with ocrelizumab. All steps of sample procurement, handling, PBMC isolation (by density centrifugation using Ficoll; GE Healthcare) cryopreservation, and subsequent thawing followed the identical standard operating procedures as previously described^20–22^.

### Immunophenotyping with intracellular staining

Human PBMC samples (1x10^7^/each) stored in liquid nitrogen were quickly thawed in a 37°C water bath and added dropwise to 10ml of 37°C pre-warmed complete RPMI (RPMI with 2mM glutamine, 1x antibiotics, 10% heat-inactivated human AB serum (Sigma H4522)) with 500 units of benzonase^®^ (Sigma E1014). After Incubation at room temperature for 5-10min, PBMCs were centrifuged at 300xg for 10min at room temperature. After removing supernatant, cells were gently resuspended in another 10ml of complete RPMI with 250 units of benzonase^®^ and centrifuged again at 300xg for 10 min at room temperature. After removing the supernatant, cells were gently resuspended in complete RPMI with 10% of heat-inactivated autologous plasma. Cells were then aliquoted, blocked with FcR blocker and stained with fluorochrome-conjugated antibodies to cell surface markers, fixed with eBioscience™ Foxp3 / Transcription Factor Staining Buffer Set (Fisher 50-1120-8857) according to manufacturer recommended protocol, then stained with mAbs to FoxP3 or other intracellularly expressed proteins. Details of antibodies, antibody panels, gating strategies and markers used to define major immune cell subsets are listed in supplementary material and method (Table 2, Supplemental Table 1, Supplemental Figure 1 and 4). The stained cells were run on Cytek® Aurora System and data were analyzed using flowjo_10.10 (BD, La Jolla, CA, USA).

**Table 2.**
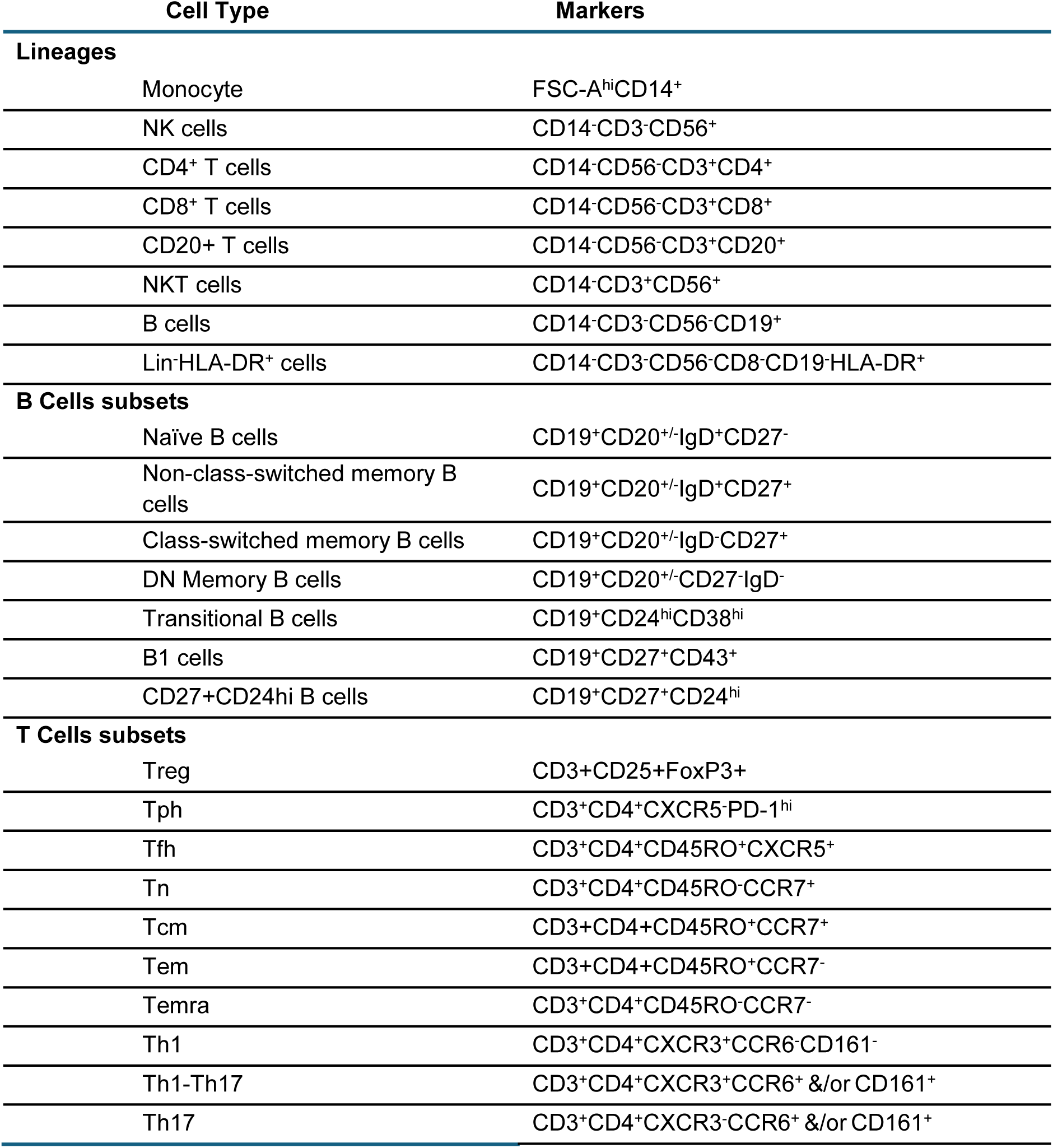
Cell lineage markers.

#### Ex vivo B cell functional assays

For ex vivo B cell cultures, PBMCs were washed 1x with complete RPMI with 10% fetal bovine serum, cultured on a 48-well flat bottom culture plate. Cells were added 400ml/well at 2.5x 10^6^/ml with or without 10μg/ml Jackson Immuno Research lab’s affinity purified F(ab’)_2_ Fragment Goat Anti-Human IgG+IgM (H+L) (Fisher NC9498449). The cells were then cultured for 48hrs at 37°C with 5%CO_2_. 1mg/ml of brefeldin A were added for last 4hrs of culture. The cells were then harvested and analyzed using flow cytometry as described earlier. The expression of cell surface molecules and intracellular cytokines were analyzed under both unstimulated and anti-IgM/IgG stimulated conditions. The differences in response to BCR-induced stimulation were evaluated using the ratio of positive frequencies under stimulated conditions (aIgM/G) over unstimulated conditions (Unst).

#### B cell subset sorting and bulk RNA-Seq

Human healthy donor PBMCs were isolated, and B cells were enriched using CD19-microbeads before downstream FACS sorting. The CD19-enriched B cells initially were gated on viable singlets. After gating out the remaining T using anti-CD3, we sorted CD24^+^CD38^+^ TransB cells and B1 subset CD43^+^CD27^+^. Naïve (CD27^+^IgD^-^) and DN (CD27^-^IgD^-^) subsets were sorted and as previously described^23^. Total RNA from sorted B cell subsets were isolated using the RNeasy Mini kit (Qiagen) with quality monitored with Agilent Bioanalyzer. Libraries were generated from polyadenylated mRNA and single-end short-read sequencing was performed (Illumina HiSeq). ∼1,000 cells per subsets were used for RNAseq.

#### RNA-Seq data analysis

Detailed methods were as previously described^23^. Briefly, raw fastq files were mapped to the hg19 version of the human genome using TopHat2 v.2.0.13 with the default parameters and the UCSC KnownGene reference transcriptome^24, 24^. Duplicate reads were removed with PICARD v.1.127. For all unique ENTREZ genes, the coverage at each exon was summarized using custom R and Bioconductor scripts and the GenomicRanges v.1.22.4 package and normalized to reads per kilobase per million reads (RPKM)^26^.

Genes were filtered for detection based on being expressed at greater than 3 reads per million in all samples for at least one group (i.e., TransB or B1 cells). Differential expression was assessed using DESeq2, comparing naïve vs. B1 and B1 vs. TransB. Genes were considered significantly differentially expressed with a Benjamini-Hochberg adjusted p-value < 0.05 and a fold change threshold of ± 1. To generate heat maps the RPKM values for all detected genes were quantile normalized and then select genes for plotting were z score normalized. Principal component analysis (PCA) was performed in R to visualize variance in gene expression profiles across samples using the prcomp function and plotted and using ggplot2. Genes contributing to the most variance were selected to generate the 3D plot.

Hierarchical clustering was performed using Wards method on Euclidean distances, followed by log transformation of RPKM values to standardize expression levels, constructing heatmaps using pHeatmap in R. Pathway analysis was performed on the DEGs using IPA and/or GO analysis in R. DEGs from each comparison were subjected to pathway enrichment analysis using the R package clusterProfiler. KEGG pathway enrichment was performed with the enrichKEGG function (organism = “hsa”, keyType = “SYMBOL”) and GO enrichment was performed with the enrichGO function (OrgDb = org.Hs.eg.db, keyType = “SYMBOL”, ont = “ALL”), both using Benjamini-Hochberg correction for multiple testing. Gene symbols were mapped to Entrez IDs using mapIds from the org.Hs.eg.db package as needed. A significance threshold of adjusted p-value < 0.05 was applied, and enriched GO terms were visualized using dot plots displaying the top categories for each comparison. All analyses were conducted in R.

#### Statistical analysis

Mixed-effects analysis followed by Dunnett’s test for multiple comparisons was used to measure statistical significance between pre-treatment and ocrelizumab-treated MS groups, while ordinary one-way ANOVA with Dunnett’s test was used to measure statistical significance between the MS pre-treatment group and HC, with p<0.05 considered significant. If data were not normally distributed, Mann-Whitney *U* tests were performed. All the statistical analyses were performed using GraphPad Prism 10.3 software (GraphPad Software, Inc. La Jolla, CA, USA).

## Results

### Ocrelizumab depletes B cells, but not CD4^+^ or CD8^+^ lymphocytes

Our prospective longitudinal study followed 36 patients before ocrelizumab and post treatment at 1-3 months, 4-7 months, 11-14 and >18 months prior to the following infusions given once every 6-7 months (Table 1). Immediate and short-term effects were reflected by the results from 1-3 months timepoints. Long-term effects were followed up beyond 18 months up to 4 years. Changes in peripheral lymphocyte populations were examined using flow cytometry on PBMCs from healthy controls (HC) and patients undergoing treatment. The frequency of B cells within PBMCs for MS patients pre-ocrelizumab treatment were 4.954±3.449% (mean ± SD) which was lower than our healthy control group (7.281±2.392, P=0.0357). A significant reduction in both the frequency (Figure 1A) and absolute number (Figure 1B) of CD19^+^ B cells within PBMCs was observed beyond 18 months after infusion, in agreement with a previous study on the anti-CD20 therapeutic rituximab^27^. A close to 90% reduction of the absolute B cell count was observed from pre-ocrelizumab (119.8±95.85 per μl) to 1-3 months (5.42±13.68, per ml) after commencement of ocrelizumab treatment (P = 0.0045) and remained low at 4-7 months (4.035±6.46, per μl)11-14 months (10.59±21.59, per μl) when measured post ocrelizumab infusions (Figure 1B). Ocrelizumab treatment did not affect the frequency or the absolute number of CD8^-^CD4^+^ (Figure 1B-C) or CD4^-^CD8^+^ T cells (Figure 1E-F) with no significant change in the CD4/CD8 T cell ratio (Figure 1G). The frequency of monocytes, NK cells, NKT or Lin^-^HLA-DR^+^ dendritic cell lineages did not change significantly with ocrelizumab (supplementary Figure 1 & 2).

**Figure 1.**
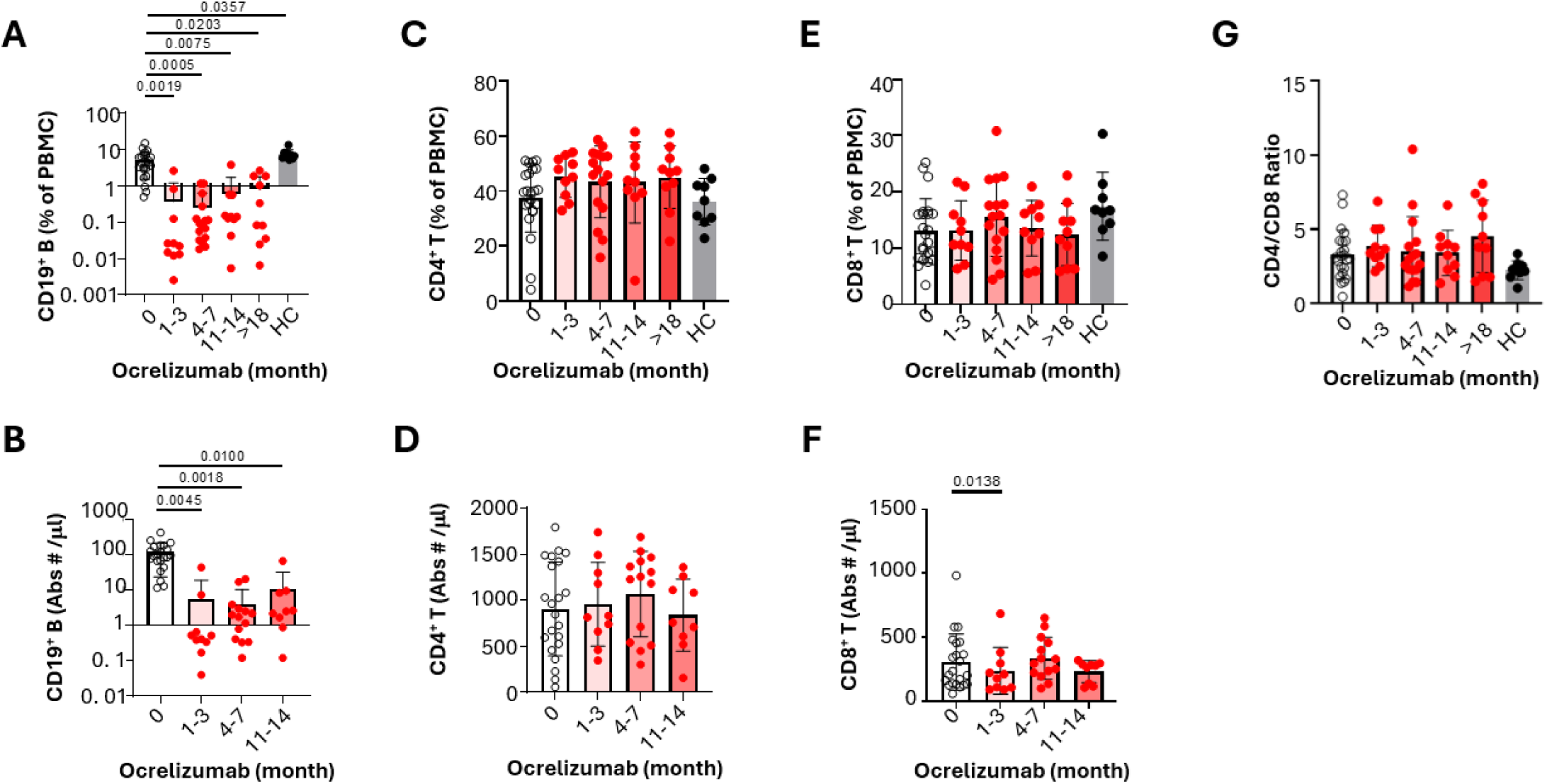
Ocrelizumab depletes B lymphocytes but not CD4^+^ or CD8^+^ T cells. To determine the short- and long-term effects of ocrelizumab on peripheral B- and T-cell populations, we assessed changes in both frequency and absolute numbers of CD19^+^ B cells, CD8^-^CD4^+^ T cells, and CD4^-^CD8^+^ T cells at baseline (0 months, pre-treatment), 1–3 months, 4–7 months, 11–14 months, and beyond 18 months post-treatment. **(A–B)** Ocrelizumab treatment led to a robust reduction in CD19^+^ B-cell frequencies **(A)** within PBMCs and absolute numbers **(B)** at all time points, and the untreated MS group was also lower than healthy controls. **(C–D)** Ocrelizumab did not alter the percentage **(C)** or absolute numbers **(D)** of CD8^-^CD4^+^ T cells. **(E–F)** Ocrelizumab did not alter the frequency **(E)** of CD4^-^CD8^+^ T cells, but there was a slight decrease in absolute numbers at 1–3 months **(F)**. **(G)** The CD4/CD8 ratio was not significantly altered. HC: untreated healthy control. Mixed-effects analysis followed by Dunnett’s test for multiple comparisons was used to measure statistical significance between pre-treatment and ocrelizumab-treated MS groups (n=23), while ordinary one-way ANOVA with Dunnett’s test was used to measure statistical significance between the MS pre-treatment group and HC (n=9), with p<0.05 considered significant.

### Ocrelizumab decreases CD20^+^ T cell populations, and moderately reduced T peripheral helper cell (Tph), but not T follicular helper cells (Tfh)

As a rare population of CD20^+^ T cells was recently identified^28^, we investigated the impact that ocrelizumab treatment had on this population. Within the first 3 months post-treatment, the frequency of total CD20^+^ T cells within CD3^+^ T cells were significantly reduced from pre-treatment 2.582±1.967% to 0.4767±0.2673% at 1-3M (P = 0.0047 Figure 2A). The frequency of CD20^+^T cells among both CD4^+^ T and CD8^+^ T cells were reduced significantly (P=0.0145 and P=0.0098 respectively Figure 2B and 2C). This reduction was persistent, since the frequency of CD20^+^ T cells among total CD3^+^ T cells, CD4^+^ T cells and CD8^+^ T cells were still significantly reduced at >18 month (P=0.027, P=0.0110, P=0.0186, respectively, Figure 2A-C). Among CD20^+^ T cells, the CD4/CD8 ratio were close to 1 in healthy control and pre-treatment MS (0.8945±0.5131, 1.201±0.669, respectively), but with Ocrelizumab treatment, some patients showed an increase in the CD4/CD8 ratio (4.824±3.417 at 1-3M, and 6.511±5.009 at >18M) within the CD20^+^ T cell population after treatment (Figure 2D).

**Figure 2.**
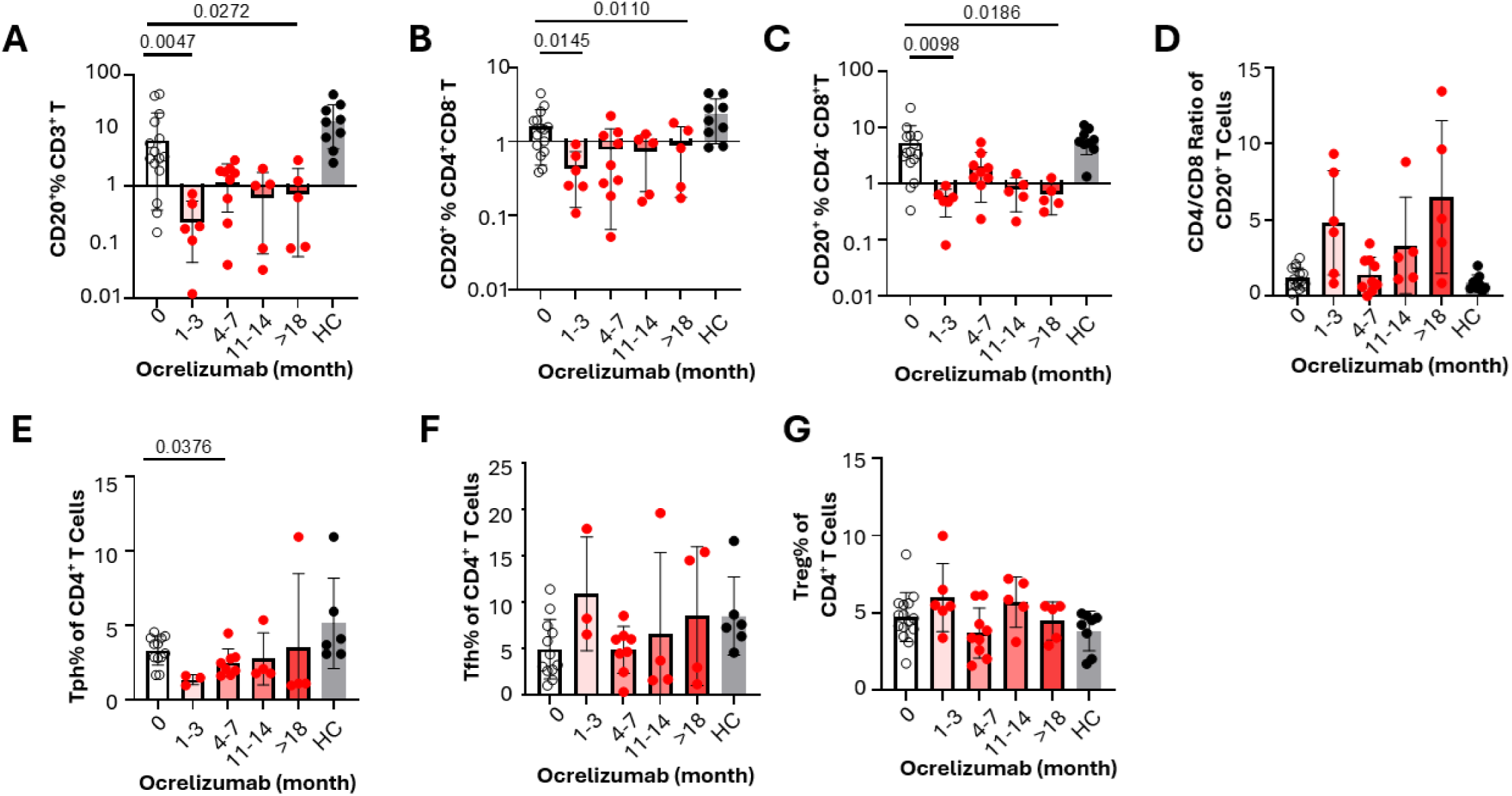
Ocrelizumab reduces CD20^+^ T cell subsets and moderately reduces Tph, but not Tfh or Treg. Effect of ocrelizumab on the frequencies of CD20^+^ T cells. **(A–D)** Treatment with ocrelizumab significantly reduced the frequency of CD20^+^ T cells at 1–3M, including reduction of CD8^-^CD4^+^CD20^+^ and CD8^+^CD4^-^CD20^+^ subsets (p < 0.05) at 1–3M and >18M. The CD4/CD8 ratio among CD20^+^ T cells **(D)** did not change significantly with treatment. Effect of ocrelizumab on the frequencies of T functional subsets. **(E–G)** Treatment with ocrelizumab significantly altered the frequencies of Tph cells (CD3^+^CD4^+^CXCR5^-^CD45RO^+^PD-1^+^) at 4–7M **(E)**. The frequencies of follicular helper T cells (CD3^+^CD4^+^CXCR5^+^) within CD3^+^CD4^+^ T cells were not significantly changed after ocrelizumab treatment (F). No significant effect of ocrelizumab was found on the frequencies of regulatory T cells (FoxP3^+^CD25^+^) within CD4^+^ T cells **(G).** Mixed-effects analysis followed by Dunnett’s test for multiple comparisons was used to measure statistical significance between pre-treatment and ocrelizumab-treated MS groups, while ordinary one-way ANOVA with Dunnett’s test for multiple comparisons was used to measure statistical significance between the MS pre-treatment group and HC with p<0.05 considered significant. Sample size **(A–D, G):** 0M: n=15, 1–3M: n=6; 4–7M: n=9; 11–14M: n=5; >18M: n=5; HC: n=9. Sample size **(E–F):** 0M: n=15, 1–3M: n=3; 4–7M: n=8; 11–14M: n=4; >18M: n=4; HC: n=6.

We further examined the changes of T cell subsets including Tn, Tcm, Tem and Temra as well as T helper subsets. Th17 and CD8 Tem were found to be reduced with ocrelizumab treatment (supplemental Fig. 3) and other subsets were not significantly changed. A role for peripheral (Tph) and follicular helper (Tfh) CD4^+^ T cells in autoimmune diseases including RA, SLE, and SS has recently been highlighted^29, 29, 31, 31^. Our data showed that the frequency of Tph (CD3^+^CD4^+^CXCR5^-^PD-1^hi^) cells were significantly reduced at 4-7 months post ocrelizumab treatment (P=0.0376, Figure 2E and Supplementary Figure 1). There were no changes in the frequency of Tfh (CD4^+^CD45RO^+^CXCR5^+^) (Figure 2F). Regulatory T cells (Treg) frequency was not significantly changed (Figure 2G).

### The selective effects of ocrelizumab on B cell subsets

To further analyze B cell subset changes, we examined residual/replenished B cell subsets with sequential gating strategies as shown in Supplemental Figure 4A. CD24^hi^CD38^hi^ Transitional B cells (TransB, also called T2-MZP B cells) - a regulatory B-cell subset -were reduced from 5.663±5.228% at 0 month to 1.155±1.1617% at 1-3 months post-treatment, but increased later over time to above 20% (Figure 3A, H). We then analyzed CD27^+^CD43^+^ B1 cells, which have innate regulatory properties. B1 showed significant resistance to ocrelizumab treatment (Figure 3B, H), with a significant increase in frequency within total B cells from 6.691±11.46% at 0 month to 63.27±17.9% at 1-3 months (P = 0.0025) and 41.71±23.37% at 4-7 months (P = 0.0094). CD27^+^ CD24^hi^ cells is another subset of B cells with potential immune regulatory function. Notably, as a fraction of total B cells, this subset was lower in the MS cohort prior to ocrelizumab (13.45±13,37%) when compared to HC (25.43±11.18%, Figure 3C; P =0.0152). After ocrelizumab treatment, the frequency of CD27^+^ CD24^hi^ cells within total B cells was reduced but not significant (Figure 3C).

**Figure 3.**
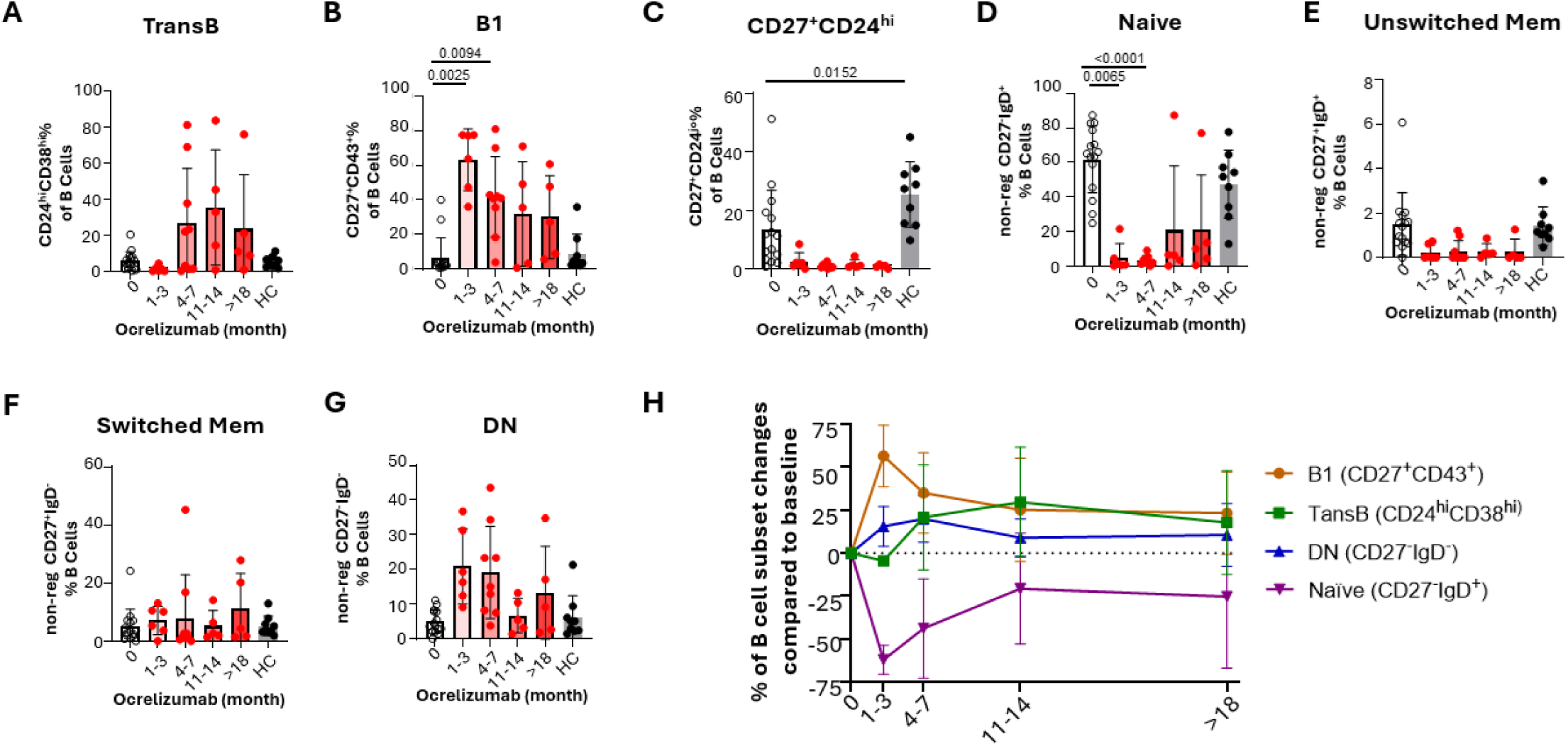
The effects of ocrelizumab on B-cell subsets. Effects of Ocrelizumab on the frequencies of various B-cell subsets within residual peripheral blood B cells were analyzed using flow cytometry and compared to pre-treatment baseline levels. **(A–C)** B-cell subsets with reported regulatory properties: The relative frequency of CD24^hi^CD38^hi^ (transitional) B cells in residual B cells showed a slight reduction at 1–3M followed by an increase at later time points, though these changes were not statistically significant **(A)**. The frequency of CD27^+^CD43^+^ (B1) cells among residual B cells was significantly increased post-treatment with significant changes at 1–3M and 4–7M **(B)**. The frequency of CD27^+^CD24hi cells among residual B cells was reduced post-treatment although the changes did not reach statistical significance, while pre-ocrelizumab MS samples showed significantly lower frequencies of this population compared to the HC group **(C)**. **(D–G)** Remaining B-cell subsets after removing those with regulatory properties: The frequency of CD27^-^IgD^+^ naïve B cells was significantly reduced at 1–3M and 4–7M **(D)**, while the frequencies of other subsets including CD27^+^IgD^+^ unswitched memory B cells **(E)**, CD27^+^IgD^-^ switched memory B cells **(F)**, and CD27^-^IgD^-^ double-negative B cells **(G)** did not show significant change post-Ocrelizumab treatment. Each time point was compared to baseline pre-treatment. Mixed-effects analysis followed by Dunnett’s test for multiple comparisons was used to measure statistical significance between pre-treatment and ocrelizumab-treated MS groups (n=15), while ordinary one-way ANOVA with Dunnett’s test for multiple comparisons was used to measure statistical significance between the MS pre-treatment group and HC (n=9), with p<0.05 considered significant. **(H)** Kinetics of major B-cell subset frequency changes compared to baseline within total B cells following Ocrelizumab treatment. 0M: n=15, 1–3M: n=6; 4–7M: n=9; 11–14M: n=5; >18M: n=5; HC: n=9.

After gating out the B cells with regulatory properties, the remaining B cells were analyzed using anti-CD27 and anti-IgD. The percentages of naïve B cells (Naïve, CD27^-^IgD^+^), within the total B cell population, were found to be significantly reduced from 61.87±19.34% at 0 month to 4.755±8.355% at 1-3 months (P = 0.0065) and 3.391±2.632 at 4-7 months (P <0.0001, Figure 3D), though gradually increased after 11-14 months and >18 months. Unswitched memory B-cells (USM, CD27^+^IgD^+^) constitute a low percentage of B cells (range: 0-6.07% of B cells) and their reduction over the course of the treatment (range: 0-1.26% of B cells) did not reach statistical significance (Figure 3E). The frequency of switched memory B-cells (SM, CD27^+^IgD^-^) was relatively unchanged (Figure 3F). Interestingly, double negative (DN, CD27^-^IgD^-^, Figure 3G) B-cells showed a trend toward increasing as compared to baseline (5.006±3.345% of B cells) at earlier time points 1-3 months (20.93±10.87% of B cells, P=0.1011) and 4-7 months (19.19±13.25% of B cells, P=0.548), and then stayed low throughout the remainder of treatment (Figure 3G and H), suggesting less efficient depletion versus other B cell subsets. To explore the reason why ocrelizumab depleted B1 and CD27^-^IgD^-^ subsets less efficiently, we assessed the CD20 expression level of all the B cell subsets that we analyzed (Supplementary Figure 4B). CD20 mean fluorescence intensity (MFI) of B1 (16988±16913, P=0.0004) and DN CD27^-^IgD^-^ (45441±52166, P=0.00162) were significantly lower than that of total B cells (76970+53904, P=0.0162), while CD24^hi^CD38^hi^ Trans B (115990±69883, P=0.0023), CD27^+^CD24^hi^ regulatory B (94036±61224, P=0.0068), and CD27^+^IgD^+^ USM (113561±62155, P=0.0015) were significantly higher than those of total B cells. Together, this data demonstrates that ocrelizumab has differential effects on B-cell subsets. The most significantly changed B cell subsets are B1, transitional B, naïve and DN, and these are shown as a kinetics diagram in Figure 3H. B1 cells are most abundant at 1-3 months, and transitional B cells are replenished first at 4-7 months post-infusion. Long term effects at 11-14 months and >18 months can also be seen as the proportions of B1 and TransB stayed higher than those of pre-treatment, while the naïve B cell population stayed lower.

### Transcriptomic analysis reveals subsets of B cell gene signatures

We sorted CD27^+^CD43^+^ B1, CD24^hi^CD38^hi^ TransB, CD19^+^IgD^+^CD27^−^Naïve, and CD19^+^IgD^−^CD27^−^DN B cells and performed RNA sequencing analysis. Principal component analysis (PCA) revealed separation of these populations (Figure 4A). Unsupervised hierarchical clustering based on variance of gene expression profiles reinforced the PCA results, grouping samples into distinct clusters that reflected their specific cell types (Figures 4B-C). We performed differential gene expression analyses to elucidate the transcriptional changes between specific cell types, comparing naïve cells to B1 cells (Figure 4D) and TransB cells to B1 cells (Supplemental Figure 5). Comparison of naïve and B1 cells identified 169 differentially expressed genes (DEGs, Figure 4B-C). Naïve cells showed higher expression of *FCN1*, *TCL1A*, *IL4R*, *SATB1*, *BACH2*, *BCL7A*, *CD72*, *CD200*, and *FCER2 (CD23)*. Many of these, such as IL4R and FCER2, endow naïve cells with heightened sensitivity to type 2 responses (IL-4), thereby increasing their ability to present antigen to T cells and to undergo antibody class switching^33^. Factors such as *BACH2* and *SATB1* are central to germinal center formation, tolerance, and the prevention of autoimmune responses^34, 34^. *CD200* has anti-inflammatory functions via *CD200R*, which reduce M1, but enhance M2-macrophages, and regulate Tregs and ultimately IL-10 production^36^, all crucial in the determination of B cell fate and the emergence of autoimmunity. Conversely, B1 cells exhibited significantly higher expression of *CD27*, *LGALS1*, *KCNN4*, *ITGB1*, *AIM2* and *IL2RB*, which can exert immunomodulatory effects, potentially promoting T cell apoptosis or skewing towards regulatory T cell (Treg) or Th2 responses, thus having a context-dependent anti-inflammatory role^37, 37^.

**Figure 4.**
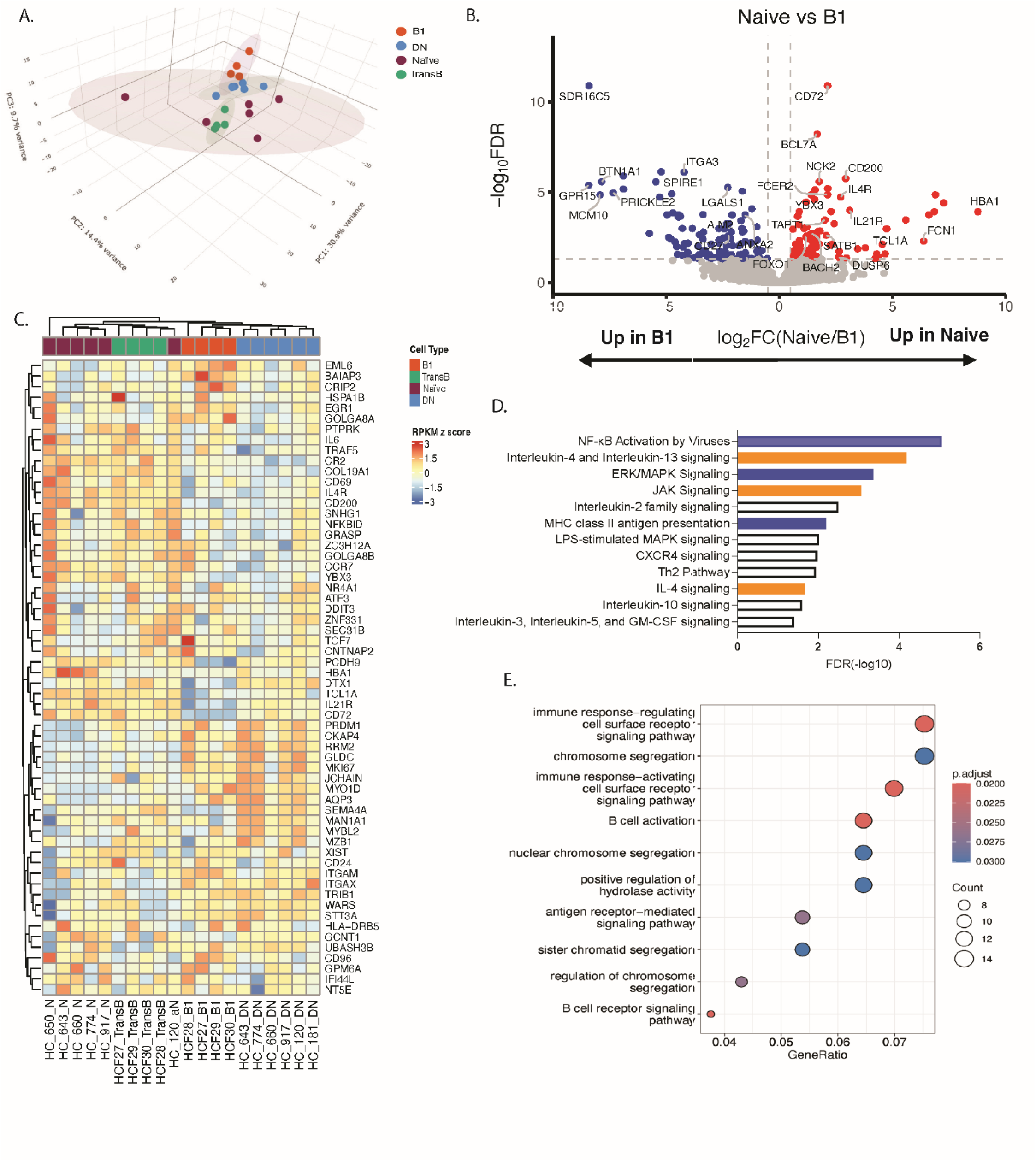
RNA-seq analysis of transitional B, B1, naïve, and DN B cells. **(A)** Principal component analysis (PCA) of transitional B, B1, naïve, and DN B cells. Each dot represents a sample. Each cell type is represented by a point and the 99% confidence interval for each cell subset is denoted by an ellipse. **(B)** Volcano plot of differentially expressed genes (FDR < 0.05, log2FC ≥ ± 0.5) between Naïve (n = 6) and B1 (n = 4) cells (top). Each point denotes a gene expressed in the RNA-seq data. **(C)** Heatmap displaying the expression of the 60 genes with the highest variance across all samples. Gene expression values were z-score normalized across rows; red indicates higher and blue indicates lower relative expression**. (D)** Significant pathways obtained from Ingenuity Pathway Analysis (IPA) of the 169 DEGs in Naive vs B1. Orange depicts predicted activation; blue indicates predicted inhibition when naïve B cells were compared to B1. **(E)** Gene Ontology (GO) enrichment analysis of DEGs from the Naïve vs B1 comparison. The dot plot displays enriched biological process terms. The gene ratio indicates the proportion of DEGs within each GO term. The size of the dot is proportional to the number of genes in the term, and the color gradient represents the adjusted p value.

While both TransB and B1 cells are defined as potentially regulatory due to their IL10 production and role in mediating tolerance, the two populations may differ in terms of tissue localization, differentiation, and functions. Comparison of TransB and B1 cells identified 131 DEGs (Supplemental Figure 5A-B). TransB cells had higher expression of IL10 signaling pathway components with increased *TCL1A*, *IL4R*, *SATB1*, *BACH2*, *IL21R*, *CD200*, and *FCER2* gene expression. The increased expression of the cytokine receptors IL4R and IL21R suggests responsiveness to T cell-derived cytokines (e.g. Th2/Tfh), crucial for guiding B cell maturation, germinal center formation, and antibody production^39^. Just as in the naïve vs B1 comparison, B1 cells displayed significantly higher expression of genes including *ITGAX (CD11c)*, *S100A4*, *ITGB1*, and *CXCR3* when compared to TransB cells. The elevated expression of *ITGAX* and *ITGB1* and *CXCR3* may be important for CNS lymphocyte trafficking and tissue homing^30, 30–42^.

### Co-signaling molecule changes in residual B cells consistent with an exhausted state after ocrelizumab treatment

To understand the activation status of the residual/replenished B cells, PBMCs from pre- and post-ocrelizumab treatment along with heathy controls were incubated for 48hrs with or without BCR stimulation using anti-IgM/IgG (Fab’)2. Without stimulation, the frequencies of co-stimulatory molecules CD40^+^ (Figure 5A) and HLA-DR^+^ (Figure 5B), but not CD80^+^ or CD86^+^ (Data not shown), were found to be significantly down-regulated in ocrelizumab treated patients (Figure 5A-B). Ocrelizumab treatment also resulted in significant up-regulation of PD-1 expression (Figure 5C) and down-regulation of PD-L1 (Figure 5D) and PD-L2 (Data not shown) in cells of B lineage compared to baseline. After B cell receptors were stimulated with anti-IgM/IgG, no notable responses were detectable for the expression CD40, HLA-DR and PD-L1 on B cells derived pre- and post-ocrelizumab treated MS patients and healthy controls. However, BCR-induced stimulation did prompt substantial upregulation of PD-1 expression on B cells from pre-ocrelizumab treated MS patients and healthy controls, but not from those derived from ocrelizumab-treated MS patients. The significance of the response difference between ocrelizumab-treated patients (T) compared with untreated patients (UT) and Healthy control (HC) are shown in supplemental Figure 6A-D. The downregulation and the unresponsiveness of CD40, HLA-DR and PD-L1 expression, imply that these cells were anergic, while the upregulation of PD-1 expression is consistent with immune exhaustion nature of these B cells.

**Figure 5.**
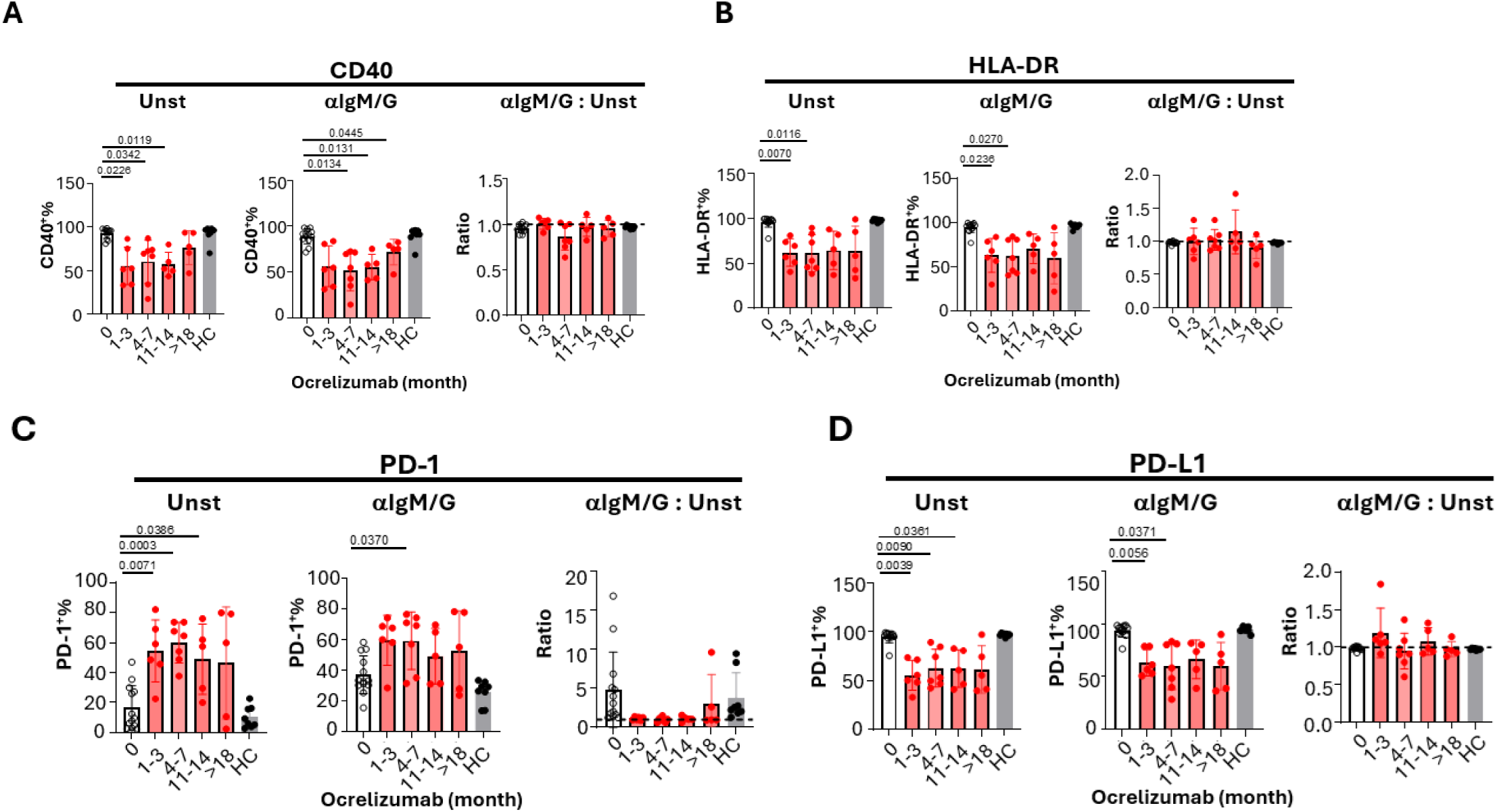
Residual B cells exhibit a dysfunctional, exhausted state. To understand the functional status of residual B cells, PBMCs from pre- and post-ocrelizumab treatment along with healthy controls were incubated for 48 hours with or without anti-IgM/IgG (Fab’)2. The frequencies of CD40^+^ **(A)**, HLA-DR^+^ **(B)**, PD-1^+^ **(C)**, and PD-L1^+^ **(D)** within B cells were analyzed using flow cytometry. For each analyte: Unst, PBMCs were unstimulated; a-IgM/G, PBMCs were stimulated with 10 μg/ml F(ab’)2 fragment of goat anti-human IgG/IgM (H+L); a-IgM/G: Unst are the ratios of the positive frequencies of analytes in B cells when stimulated with anti-IgM/G (a-IgM/G) over those without stimulation (Unst). Mixed-effects analysis followed by Dunnett’s test for multiple comparisons was used to measure statistical significance between pre-treatment and ocrelizumab-treated MS groups (n=13), while ordinary one-way ANOVA with Dunnett’s test for multiple comparisons was used to measure statistical significance between the MS pre-treatment group and HC (n=9), with p<0.05 considered significant. 0M: n=13, 1–3M: n=6; 4–7M: n=7; 11–14M: n=5; >18M: n=5; HC: n=8.

### Residual B-cells in ocrelizumab-treated patients co-expressed pro-inflammatory cytokines and are anergic with an increased IL10/TNFα and IL10/IL6 ratio

To understand the function of cytokine production by the residua/replenishing B cells, PBMCs from pre- and post-ocrelizumab treatment along with heathy controls were incubated for 48hrs with or without anti-IgM/IgG (Fab’)2. The residual/replenishing B cells derived from ocrelizumab-treated MS patients co-expressed high levels of IFNγ (Figure 6A, left-hand panel), TNFα (Figure 6B, left-hand panel), IL-6 (Figure 6C, left-hand panel), and IL-10 (Figure 6D, left-hand panel) under unstimulated condition compared with those derived from pre-ocrelizumab treated MS patients and healthy controls, but unlike those B cells from pre-Ocrelizumab MS patients and healthy controls, the level of these pro-inflammatory cytokines production in B cells from ocrelizumab-treated patients could not be further enhanced by anti-IgM/IgG stimulation (Fig. 6A-C, middle and right-hand panel Supplementary Figure 6 E-G), Corroborating with CD40 and HLA-DR, the unresponsive behavior of these cells suggests they were anergic. Unlike IFNγ, TNFα and IL-6, IL-10 expression was increased in B cell derived from ocrelizumab treated MS patients compared to pre-ocrelizumab treated MS patients and healthy controls under unstimulated condition. BCR stimulation did not alter the express of IL-10 significantly in both pre- and post-ocrelizumab-treated samples (Figure 6D, Supplementary Figure H).

**Figure 6.**
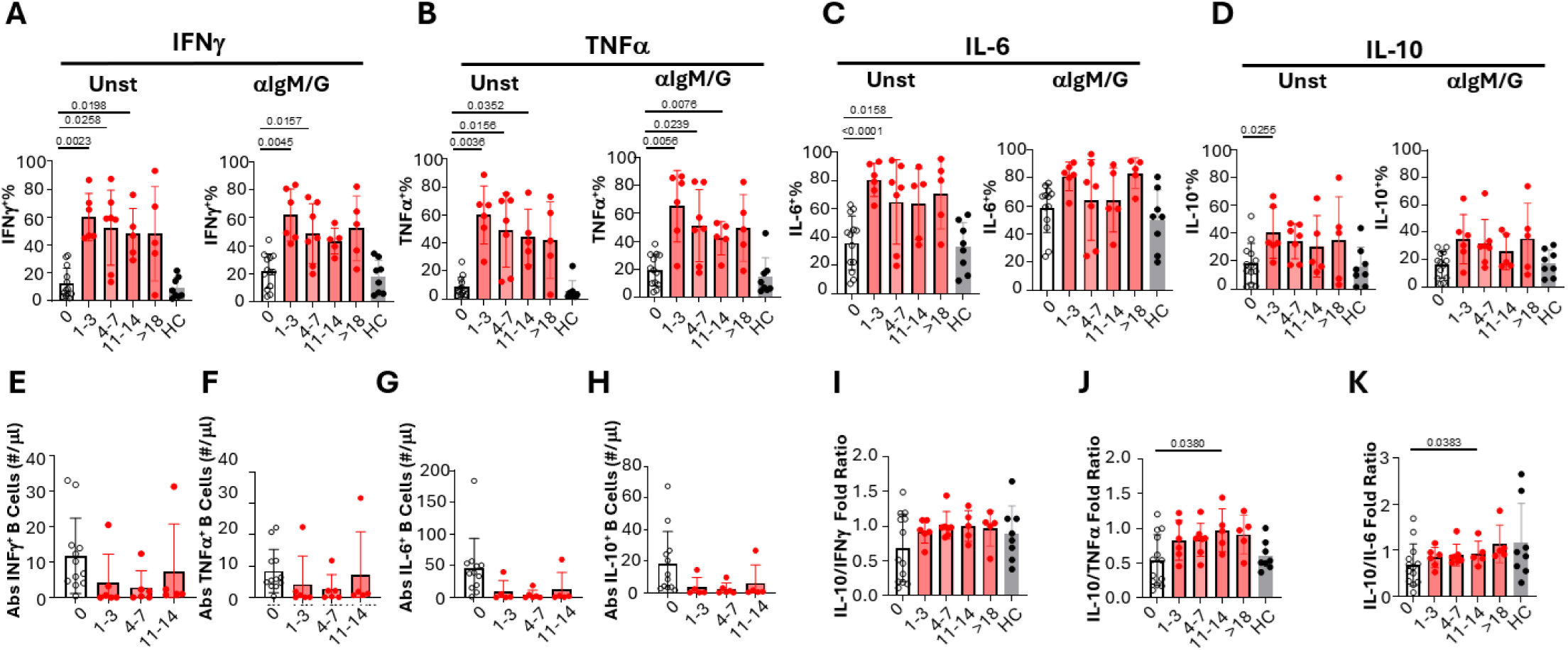
Residual B cells co-expressed pro-inflammatory cytokines but are functionally anergic with increased anti- to pro-inflammatory cytokine ratios after ocrelizumab treatment. To understand the cytokine profile of residual B cells, PBMCs from pre- and post-ocrelizumab treatment along with healthy controls were incubated for 48 hours with or without anti-IgM/IgG (Fab’)2. The frequencies of IFNγ^+^ **(A)**, TNFα^+^ **(B)**, IL-6^+^ **(C)**, and IL-10^+^ **(D)** within B cells were analyzed using flow cytometry. For each cytokine: Unst, PBMCs were unstimulated; a-IgM/G, PBMCs were stimulated with 10 μg/ml F(ab’)2 fragment of goat anti-human IgG/IgM (H+L); a-IgM/G: Unst are the ratios of the positive frequencies of analytes in B cells when stimulated with anti-IgM/G (a-IgM/G) over those without stimulation (Unst). **(E–H)** The absolute number of cytokine^+^ B cells (#/μl) was calculated using the positive frequencies of these cytokines under unstimulated conditions and the total number of B cells in blood for IFNγ **(E)**, TNFα **(F)**, IL-6 (G), and IL-10 (H). **(I–K)** The ratios of IL-10^+^ frequency change (fold) under a-IgM/G stimulation over pro-inflammatory cytokine^+^ frequency change (fold) under a-IgM/G stimulation were calculated for IFNγ **(I)**, TNFα **(J)**, and IL-6 **(K)**. Mixed-effects analysis followed by Dunnett’s test for multiple comparisons was used to measure statistical significance between pre-treatment and ocrelizumab-treated MS groups (n=13), while ordinary one-way ANOVA with Dunnett’s test for multiple comparisons was used to measure statistical significance between the MS pre-treatment group and HC (n=9), with p<0.05 considered significant. 0M: n=13, 1–3M: n=6; 4–7M: n=7; 11–14M: n=5; >18M: n=5; HC: n=8.

To evaluate the overall cytokine change in the residual/replenishing B cells, we calculated the absolute numbers of IFNγ (Figure 6E), TNFα (Figure 6F), IL-6 (Figure 6G) and IL-10-producing (Figure 6H) B cells from pre- and post-ocrelizumab treated MS patients using the frequencies of unstimulated cytokine positive B cells times the total absolute number of B cells. We found that within the limited amount of remaining B cells, the overall B cells were less inflammatory.

To analyze the overall effect of BCR-induced stimulation, we analyzed IL-10 responses compared to those responses of pro-inflammatory cytokines. We found that the anti-inflammatory IL-10 response to BCR stimulation were superior to those proinflammatory cytokines responses (Figure I-K), particularly IL-10 to TNFα response ratio (P=0.0380 Figure 6J) and IL-10 to IL-6 response ratio (P=0.0383, Figure 6K) at 11-14 months. These data are consistent with RNAseq analysis of B1 and transitional B cells showing increased IL10 pathways (Figure 4 and supplemental 5).

## Discussion

Ocrelizumab treatment reshapes the immunological landscape in MS patients, selectively depleting certain B cell populations while preserving others with key regulatory functions. Our comprehensive prospective longitudinal study of 36 MS patients reveals a mechanism of action beyond simple pan-B-cell depletion, potentially explaining the drug’s sustained therapeutic efficacy. While we confirmed the expected persistent depletion of total B cells over the 18-month treatment period, CD4^+^, CD8^+^ T cells, NK, NKT, monocyte and Lin^-^HLA-DR^+^ lineages are not significantly changed (Figure 1, Supplemental 1&2).

Our study also revealed an effect of ocrelizumab on CD20-expressing T cells, a newly recognized subpopulation involved in autoimmune disorders^43^. Ocrelizumab significantly reduced the total number of CD20^+^ T cells, including both CD4^+^CD20^+^ and CD8^+^CD20^+^ subsets (Figures 2). This presents an additional mechanism by which ocrelizumab may affect T cell-mediated pathology in MS, beyond its B cell depletion. Notably, the varying effects on T helper subsets showed a decrease in peripheral helper T cells (Tph), while follicular helper T cells (Tfh) remained stable (Figures 2). This selective reduction of Tph cells may be crucial for disrupting pathogenic T-B cell interactions in MS. The decrease in Th17 and Tem post-treatment (Supplemental 3) suggest a shift toward immune regulation^44, 44^. Tph cells orchestrate extrafollicular autoreactive B cell responses through IL-21 and have been implicated in the suppression of regulatory B cell development^29, 29, 45, 45^. Thus, their selective reduction by ocrelizumab could disrupt pathogenic B cell support while facilitating the expansion of regulatory B cell networks. Studies demonstrating Tph-mediated suppression of regulatory B cells in other inflammatory conditions support this hypothesis^22, 22, 45^.

We further observed that ocrelizumab resulted in a varying depletion across B cell subsets, revealing a sophisticated reshaping of the B cell compartment^47^. Naïve B cells (CD27^-^IgD^+^) showed the greatest reduction in response to ocrelizumab but gradually recovered and were maintained below baseline levels by 11-14 months and beyond 18 months (Figure 3D and 3H). Conversely, unswitched memory B cells (CD27^+^IgD^+^) remained persistently low (Figure 3E), while double-negative B cells (CD27^-^IgD^-^) demonstrated relative resistance to depletion (Figure 3). Importantly, our data showed that CD24^hi^CD38^hi^ TransB replenished first (Figure 3A) and gradually increased over time and were maintained above baseline (Figure 3H). CD27^+^CD43^+^ B1 cells are more abundant in the B cell compartment post-treatment at early time point 1-3 months (Figure 3B and 3H) suggesting they are resistant to depletion, possibly due to their relative lower expression level of CD20 (Supplemental Figure 4B). These findings indicate that ocrelizumab may selectively spare or potentially promote B cell subsets with immunoregulatory capacity. This differential susceptibility may reflect differences in tissue residency among these subsets.

Our previous work showed a decrease in number and/or functional capacity in two Breg subsets B1 CD27^+^CD43^+^CD19^+^ and the TransB CD38^hi^CD24^hi^CD19^+^ in MS and we were the first to show that both types of Bregs are increased in response to dimethyl fumarate (DMF) treatment^20^, which were validated by larger study ^48^. We also showed that the frequencies and function of two subsets of Breg were increased with siponimod^22^. The fact that several DMTs with different mechanisms of actions, all converge upon upregulating B1 and TransB is intriguing. We therefore sorted these populations of regulatory B cells and compared them to pathogenic B cell subsets (naïve and DN) B cells.

The transcriptional profiles of these preserved regulatory CD24^hi^CD38^hi^ TransB and CD27^+^CD43^+^ B1 cell subsets provide mechanistic insights into their potential functions. CD24^hi^CD38^hi^ TransB cells exhibited elevated expressions of IL4R, IL21R, and FCER2, indicating responsiveness to T cell-derived cytokines that promote IL-10 production (Supplemental Figure 5). The elevated expression of CD200 in TransB cells, known to induce Treg polarization, suggests potential to amplify regulatory networks^36^. In contrast, B1 cells (CD27^+^CD43^+^) displayed higher levels of IL2RB and LGALS1 (Galectin-1) (Figure 4, Supplemental Figure 5), which can induce regulatory T cells and promote an anti-inflammatory environment through both IL-10-dependent and independent mechanisms^10, 10^. Recent evidence that Tph cells suppress regulatory B cells in inflammatory conditions further supports this model^50^. The reduction in Tph cells following ocrelizumab treatment may contribute to the relative enrichment of B1 cells, altering the IL-21/IL21R axis and promoting a shift towards more anti-inflammatory states. While further investigations are needed to fully characterize these B1-Tph interactions and their impact on cytokine production, our findings suggest this axis represents a key mechanism underlying ocrelizumab’s therapeutic effects.

Ocrelizumab treatment resulted in significant down-regulation of co-signaling molecules CD40, HLA-DR, PD-L1 and PD-L2 and up-regulation of PD-1 in residual/replenished B cells. Ongoing clinical trials specifically targeting co-signaling such as anti-CD40L to intercept B-T interactions may yield safer and more efficacious immunoregulation^51–52^. Our data showing residual/replenishing of B cells with increased IFNγ, TNFα, and IL-6 secretion in the unstimulated state suggest that the residual/replenished B cells potentially have a reinforced proinflammatory capacity, which would warrant continued ocrelizumab treatment. On the other hand, the presence of residual proinflammatory cytokines may also help to fight off infection and malignancy. The residual B-cells in ocrelizumab-treated MS patients, though co-expressing pro-inflammatory cytokines, also have an exhausted phenotype. Most importantly, ex vivo BCR stimulation with anti-IgM/IgG showed the relative capacity for proinflammatory cytokine production was reduced with a significantly increased IL-10/TNFα and IL-10/IL6 ratio in the residual/replenishing B cells following ocrelizumab treatment.

Collectively, these data suggest that ocrelizumab treatment creates a self-reinforcing regulatory circuit: the reduction of Tph cells alleviates the suppression of regulatory B cells, particularly the B1 cells, which subsequently expand and further promote regulatory T cell networks through mechanisms mediated by IL2RB, LGALS1, and potentially IL-10. The decreased expression of IL21R in B1 cells may offer additional protection from Tph-mediated effects, explaining their persistence despite the overall depletion of B cells^50^. Simultaneously, TransB cells utilize their responsiveness to cytokines and the expression of CD200 to reinforce this regulatory environment. This immunological reshaping, characterized by the preservation of regulatory B cells and reduction of pathogenic T and B cells offers new insights into ocrelizumab’s mechanism of action and suggests potential approaches for developing more targeted MS therapies.

## Author Contributions

All authors participated in the interpretation of study results and in the drafting, critical revision, and approval of the final version of the manuscript. QW and YMD conceived the study. QWu, MGR, QWang, DD, CDS, JB, DED, PLC, SKL, DAF and YMD contributed to the acquisition of study results. QW, MGR, JG, JSM, CDS, DED and YMD performed the analysis of study results. QW, MGR, EAM, DAF and YMD drafted and edited the manuscript.

## Acknowledgement

We thank Sophina H Taitano, Luciën E P M van der Vlugt and U Michigan Flowcytometry Core for helping sorting B cells. We also thank OMRF Center for Biomedical Data Sciences (CBDS) for helping with RNA seq analysis. We sincerely thank all of our patients and families for participating in the study.

## REFERENCES

1. Hauser SL, Oksenberg JR. The neurobiology of multiple sclerosis: genes, inflammation, and neurodegeneration. Neuron. 2006;52(1):61–76. doi: 10.1016/j.neuron.2006.09.011. PMID: 17015227.

2. Trapp BD, Nave KA. Multiple sclerosis: an immune or neurodegenerative disorder? Annu Rev Neurosci. 2008;31:247–69. doi: 10.1146/annurev.neuro.30.051606.094313. PMID: 18558855.

3. Bar-Or A, Pachner A, Menguy-Vacheron F, Kaplan J, Wiendl H. Teriflunomide and its mechanism of action in multiple sclerosis. Drugs. 2014;74(6):659–74. doi: 10.1007/s40265-014-0212-x. PMID: 24740824; PMCID: PMC4003395.

4. Du FH, Mills EA, Mao-Draayer Y. Next-generation anti-CD20 monoclonal antibodies in autoimmune disease treatment. Auto Immun Highlights. 2017;8(1):12. Epub 20171116. doi: 10.1007/s13317-017-0100-y. PMID: 29143151; PMCID: PMC5688039.

5. Hausler D, Hausser-Kinzel S, Feldmann L, Torke S, Lepennetier G, Bernard CCA, Zamvil SS, Bruck W, Lehmann-Horn K, Weber MS. Functional characterization of reappearing B cells after anti-CD20 treatment of CNS autoimmune disease. Proc Natl Acad Sci U S A. 2018;115(39):9773–8. Epub 20180907. doi: 10.1073/pnas.1810470115. PMID: 30194232; PMCID: PMC6166805.

6. Lehmann-Horn K, Kronsbein HC, Weber MS. Targeting B cells in the treatment of multiple sclerosis: recent advances and remaining challenges. Ther Adv Neurol Disord. 2013;6(3):161–73. doi: 10.1177/1756285612474333. PMID: 23634189; PMCID: PMC3625013.

7. Li R, Rezk A, Miyazaki Y, Hilgenberg E, Touil H, Shen P, Moore CS, Michel L, Althekair F, Rajasekharan S, Gommerman JL, Prat A, Fillatreau S, Bar-Or A, Canadian BciMST. Proinflammatory GM-CSF-producing B cells in multiple sclerosis and B cell depletion therapy. Sci Transl Med. 2015;7(310):310ra166. doi: 10.1126/scitranslmed.aab4176. PMID: 26491076.

8. Blair PA, Norena LY, Flores-Borja F, Rawlings DJ, Isenberg DA, Ehrenstein MR, Mauri C. CD19(+)CD24(hi)CD38(hi) B cells exhibit regulatory capacity in healthy individuals but are functionally impaired in systemic Lupus Erythematosus patients. Immunity. 2010;32(1):129–40. Epub 20100114. doi: 10.1016/j.immuni.2009.11.009. PMID: 20079667.

9. Iwata Y, Matsushita T, Horikawa M, Dilillo DJ, Yanaba K, Venturi GM, Szabolcs PM, Bernstein SH, Magro CM, Williams AD, Hall RP, St Clair EW, Tedder TF. Characterization of a rare IL-10-competent B-cell subset in humans that parallels mouse regulatory B10 cells. Blood. 2011;117(2):530–41. Epub 20101020. doi: 10.1182/blood-2010-07-294249. PMID: 20962324; PMCID: PMC3031478.

10. Griffin DO, Rothstein TL. Human “orchestrator” CD11b(+) B1 cells spontaneously secrete interleukin-10 and regulate T-cell activity. Mol Med. 2012;18(1):1003–8. Epub 20120907. doi: 10.2119/molmed.2012.00203. PMID: 22634719; PMCID: PMC3459484.

11. van de Veen W, Stanic B, Wirz OF, Jansen K, Globinska A, Akdis M. Role of regulatory B cells in immune tolerance to allergens and beyond. J Allergy Clin Immunol. 2016;138(3):654–65. doi: 10.1016/j.jaci.2016.07.006. PMID: 27596706.

12. de Masson A, Bouaziz JD, Le Buanec H, Robin M, O’Meara A, Parquet N, Rybojad M, Hau E, Monfort JB, Branchtein M, Michonneau D, Dessirier V, Sicre de Fontbrune F, Bergeron A, Itzykson R, Dhedin N, Bengoufa D, Peffault de Latour R, Xhaard A, Bagot M, Bensussan A, Socie G. CD24(hi)CD27(+) and plasmablast-like regulatory B cells in human chronic graft-versus-host disease. Blood. 2015;125(11):1830–9. Epub 20150120. doi: 10.1182/blood-2014-09-599159. PMID: 25605369.

13. Hasan MM, Thompson-Snipes L, Klintmalm G, Demetris AJ, O’Leary J, Oh S, Joo H. CD24(hi)CD38(hi) and CD24(hi)CD27(+) Human Regulatory B Cells Display Common and Distinct Functional Characteristics. J Immunol. 2019;203(8):2110–20. Epub 20190911. doi: 10.4049/jimmunol.1900488. PMID: 31511354.

14. Kantor AB, Stall AM, Adams S, Herzenberg LA, Herzenberg LA. Differential development of progenitor activity for three B-cell lineages. Proc Natl Acad Sci U S A. 1992;89(8):3320–4. doi: 10.1073/pnas.89.8.3320. PMID: 1565622; PMCID: PMC48858.

15. Ghosn EE, Sadate-Ngatchou P, Yang Y, Herzenberg LA, Herzenberg LA. Distinct progenitors for B-1 and B-2 cells are present in adult mouse spleen. Proc Natl Acad Sci U S A. 2011;108(7):2879–84. Epub 20110131. doi: 10.1073/pnas.1019764108. PMID: 21282663; PMCID: PMC3041118.

16. Choi YS, Dieter JA, Rothaeusler K, Luo Z, Baumgarth N. B-1 cells in the bone marrow are a significant source of natural IgM. Eur J Immunol. 2012;42(1):120–9. Epub 20111128. doi: 10.1002/eji.201141890. PMID: 22009734; PMCID: PMC3426357.

17. Flores-Borja F, Bosma A, Ng D, Reddy V, Ehrenstein MR, Isenberg DA, Mauri C. CD19+CD24hiCD38hi B cells maintain regulatory T cells while limiting TH1 and TH17 differentiation. Sci Transl Med. 2013;5(173):173ra23. doi: 10.1126/scitranslmed.3005407. PMID: 23427243.

18. Lampropoulou V, Hoehlig K, Roch T, Neves P, Calderon Gomez E, Sweenie CH, Hao Y, Freitas AA, Steinhoff U, Anderton SM, Fillatreau S. TLR-activated B cells suppress T cell-mediated autoimmunity. J Immunol. 2008;180(7):4763–73. doi: 10.4049/jimmunol.180.7.4763. PMID: 18354200.

19. Pennati A, Ng S, Wu Y, Murphy JR, Deng J, Rangaraju S, Asress S, Blanchfield JL, Evavold B, Galipeau J. Regulatory B Cells Induce Formation of IL-10-Expressing T Cells in Mice with Autoimmune Neuroinflammation. J Neurosci. 2016;36(50):12598–610. Epub 20161107. doi: 10.1523/JNEUROSCI.1994-16.2016. PMID: 27821578; PMCID: PMC5157105.

20. Lundy SK, Wu Q, Wang Q, Dowling CA, Taitano SH, Mao G, Mao-Draayer Y. Dimethyl fumarate treatment of relapsing-remitting multiple sclerosis influences B-cell subsets. Neurol Neuroimmunol Neuroinflamm. 2016;3(2):e211. Epub 20160303. doi: 10.1212/NXI.0000000000000211. PMID: 27006972; PMCID: PMC4784801.

21. Wu Q, Wang Q, Mao G, Dowling CA, Lundy SK, Mao-Draayer Y. Dimethyl Fumarate Selectively Reduces Memory T Cells and Shifts the Balance between Th1/Th17 and Th2 in Multiple Sclerosis Patients. J Immunol. 2017;198(8):3069–80. Epub 20170303. doi: 10.4049/jimmunol.1601532. PMID: 28258191; PMCID: PMC5464403.

22. Wu Q, Mills EA, Wang Q, Dowling CA, Fisher C, Kirch B, Lundy SK, Fox DA, Mao-Draayer Y, Group AMSS. Siponimod enriches regulatory T and B lymphocytes in secondary progressive multiple sclerosis. JCI Insight. 2020;5(3). Epub 20200213. doi: 10.1172/jci.insight.134251. PMID: 31935197; PMCID: PMC7098784.

23. Scharer CD, Blalock EL, Mi T, Barwick BG, Jenks SA, Deguchi T, Cashman KS, Neary BE, Patterson DG, Hicks SL, Khosroshahi A, Eun-Hyung Lee F, Wei C, Sanz I, Boss JM. Epigenetic programming underpins B cell dysfunction in human SLE. Nat Immunol. 2019;20(8):1071–82. Epub 20190701. doi: 10.1038/s41590-019-0419-9. PMID: 31263277; PMCID: PMC6642679.

24. Kim D, Pertea G, Trapnell C, Pimentel H, Kelley R, Salzberg SL. TopHat2: accurate alignment of transcriptomes in the presence of insertions, deletions and gene fusions. Genome Biol. 2013;14(4):R36. Epub 20130425. doi: 10.1186/gb-2013-14-4-r36. PMID: 23618408; PMCID: PMC4053844.

25. Kuhn RM, Haussler D, Kent WJ. The UCSC genome browser and associated tools. Brief Bioinform. 2013;14(2):144–61. Epub 20120820. doi: 10.1093/bib/bbs038. PMID: 22908213; PMCID: PMC3603215.

26. Lawrence M, Huber W, Pages H, Aboyoun P, Carlson M, Gentleman R, Morgan MT, Carey VJ. Software for computing and annotating genomic ranges. PLoS Comput Biol. 2013;9(8):e1003118. Epub 20130808. doi: 10.1371/journal.pcbi.1003118. PMID: 23950696; PMCID: PMC3738458.

27. Nissimov N, Hajiyeva Z, Torke S, Grondey K, Bruck W, Hausser-Kinzel S, Weber MS. B cells reappear less mature and more activated after their anti-CD20-mediated depletion in multiple sclerosis. Proc Natl Acad Sci U S A. 2020;117(41):25690–9. Epub 20200930. doi: 10.1073/pnas.2012249117. PMID: 32999069; PMCID: PMC7568262.

28. Shinoda K, Li R, Rezk A, Mexhitaj I, Patterson KR, Kakara M, Zuroff L, Bennett JL, von Budingen HC, Carruthers R, Edwards KR, Fallis R, Giacomini PS, Greenberg BM, Hafler DA, Ionete C, Kaunzner UW, Lock CB, Longbrake EE, Pardo G, Piehl F, Weber MS, Ziemssen T, Jacobs D, Gelfand JM, Cross AH, Cameron B, Musch B, Winger RC, Jia X, Harp CT, Herman A, Bar-Or A. Differential effects of anti-CD20 therapy on CD4 and CD8 T cells and implication of CD20-expressing CD8 T cells in MS disease activity. Proc Natl Acad Sci U S A. 2023;120(3):e2207291120. Epub 20230112. doi: 10.1073/pnas.2207291120. PMID: 36634138; PMCID: PMC9934304.

29. Yoshitomi H, Ueno H. Shared and distinct roles of T peripheral helper and T follicular helper cells in human diseases. Cell Mol Immunol. 2021;18(3):523–7. Epub 20200831. doi: 10.1038/s41423-020-00529-z. PMID: 32868910; PMCID: PMC8027819.

30. Bocharnikov AV, Keegan J, Wacleche VS, Cao Y, Fonseka CY, Wang G, Muise ES, Zhang KX, Arazi A, Keras G, Li ZJ, Qu Y, Gurish MF, Petri M, Buyon JP, Putterman C, Wofsy D, James JA, Guthridge JM, Diamond B, Anolik JH, Mackey MF, Alves SE, Nigrovic PA, Costenbader KH, Brenner MB, Lederer JA, Rao DA, Accelerating Medicines Partnership RASLEN. PD-1hiCXCR5-T peripheral helper cells promote B cell responses in lupus via MAF and IL-21. JCI Insight. 2019;4(20). Epub 20191017. doi: 10.1172/jci.insight.130062. PMID: 31536480; PMCID: PMC6824311.

31. Zhu Y, Jiang Q, Lei C, Yu Q, Qiu L. The response of CD27(+)CD38(+) plasmablasts, CD24(hi)CD38(hi) transitional B cells, CXCR5(-)ICOS(+)PD-1(+) Tph, Tph2 and Tfh2 subtypes to allergens in children with allergic asthma. BMC Pediatr. 2024;24(1):154. Epub 20240229. doi: 10.1186/s12887-024-04622-4. PMID: 38424520; PMCID: PMC10902953.

32. Schmitt N, Bentebibel SE, Ueno H. Phenotype and functions of memory Tfh cells in human blood. Trends Immunol. 2014;35(9):436–42. Epub 20140703. doi: 10.1016/j.it.2014.06.002. PMID: 24998903; PMCID: PMC4152409.

33. Aranda CJ, Gonzalez-Kozlova E, Saunders SP, Fernandes-Braga W, Ota M, Narayanan S, He JS, Del Duca E, Swaroop B, Gnjatic S, Shattner G, Reibman J, Soter NA, Guttman-Yassky E, Curotto de Lafaille MA. IgG memory B cells expressing IL4R and FCER2 are associated with atopic diseases. Allergy. 2023;78(3):752–66. Epub 20221219. doi: 10.1111/all.15601. PMID: 36445014; PMCID: PMC9991991.

34. Hu Q, Xu T, Zhang W, Huang C. Bach2 regulates B cell survival to maintain germinal centers and promote B cell memory. Biochem Biophys Res Commun. 2022;618:86–92. Epub 20220608. doi: 10.1016/j.bbrc.2022.06.009. PMID: 35716600.

35. Ozawa T, Fujii K, Sudo T, Doi Y, Nakai R, Shingai Y, Ueda T, Baba Y, Hosen N, Yokota T. Special AT-Rich Sequence-Binding Protein 1 Supports Survival and Maturation of Naive B Cells Stimulated by B Cell Receptors. J Immunol. 2022;208(8):1937–46. Epub 20220404. doi: 10.4049/jimmunol.2101097. PMID: 35379742.

36. Chu KH, Chiang BL. CD200R activation on naive T cells by B cells induces suppressive activity of T cells via IL-24. Cell Mol Life Sci. 2024;81(1):231. Epub 20240523. doi: 10.1007/s00018-024-05268-2. PMID: 38780647; PMCID: PMC11116298.

37. Tumang JR, Holodick NE, Vizconde TC, Kaku H, Frances R, Rothstein TL. A CD25(-) positive population of activated B1 cells expresses LIFR and responds to LIF. Front Immunol. 2011;2:6. Epub 20110321. doi: 10.3389/fimmu.2011.00006. PMID: 22566797; PMCID: PMC3342026.

38. Yu X, Qian J, Ding L, Yin S, Zhou L, Zheng S. Galectin-1: A Traditionally Immunosuppressive Protein Displays Context-Dependent Capacities. Int J Mol Sci. 2023;24(7). Epub 20230330. doi: 10.3390/ijms24076501. PMID: 37047471; PMCID: PMC10095249.

39. Shehata L, Thouvenel CD, Hondowicz BD, Pew LA, Pritchard GH, Rawlings DJ, Choi J, Pepper M. Interleukin-4 downregulates transcription factor BCL6 to promote memory B cell selection in germinal centers. Immunity. 2024;57(4):843–58 e5. Epub 20240320. doi: 10.1016/j.immuni.2024.02.018. PMID: 38513666; PMCID: PMC11104266.

40. Cheekatla SS, Tripathi D, Venkatasubramanian S, Paidipally P, Welch E, Tvinnereim AR, Nurieva R, Vankayalapati R. IL-21 Receptor Signaling Is Essential for Optimal CD4(+) T Cell Function and Control of Mycobacterium tuberculosis Infection in Mice. J Immunol. 2017;199(8):2815–22. Epub 20170830. doi: 10.4049/jimmunol.1601231. PMID: 28855309; PMCID: PMC5636679.

41. Jenks SA, Cashman KS, Zumaquero E, Marigorta UM, Patel AV, Wang X, Tomar D, Woodruff MC, Simon Z, Bugrovsky R, Blalock EL, Scharer CD, Tipton CM, Wei C, Lim SS, Petri M, Niewold TB, Anolik JH, Gibson G, Lee FE, Boss JM, Lund FE, Sanz I. Distinct Effector B Cells Induced by Unregulated Toll-like Receptor 7 Contribute to Pathogenic Responses in Systemic Lupus Erythematosus. Immunity. 2018;49(4):725–39 e6. Epub 20181009. doi: 10.1016/j.immuni.2018.08.015. PMID: 30314758; PMCID: PMC6217820.

42. Akama-Garren EH, Carroll MC. T cell help in the autoreactive germinal center. Scand J Immunol. 2022;95(6):e13192. Epub 20220531. doi: 10.1111/sji.13192. PMID: 35587582; PMCID: PMC9250425.

43. Vlaming M, Bilemjian V, Freile JA, Lourens HJ, van Rooij N, Huls G, van Meerten T, de Bruyn M, Bremer E. CD20 positive CD8 T cells are a unique and transcriptionally-distinct subset of T cells with distinct transmigration properties. Sci Rep. 2021;11(1):20499. Epub 20211015. doi: 10.1038/s41598-021-00007-0. PMID: 34654826; PMCID: PMC8520003.

44. Yang X, Yang J, Chu Y, Xue Y, Xuan D, Zheng S, Zou H. T follicular helper cells and regulatory B cells dynamics in systemic lupus erythematosus. PLoS One. 2014;9(2):e88441. Epub 20140214. doi: 10.1371/journal.pone.0088441. PMID: 24551101; PMCID: PMC3925141.

45. Ding T, Su R, Wu R, Xue H, Wang Y, Su R, Gao C, Li X, Wang C. Frontiers of Autoantibodies in Autoimmune Disorders: Crosstalk Between Tfh/Tfr and Regulatory B Cells. Front Immunol. 2021;12:641013. Epub 20210326. doi: 10.3389/fimmu.2021.641013. PMID: 33841422; PMCID: PMC8033031.

46. Fazazi MR, Doss P, Pereira R, Fudge N, Regmi A, Joly-Beauparlant C, Akbar I, Yeola AP, Mailhot B, Baillargeon J, Grenier P, Bertrand N, Lacroix S, Droit A, Moore CS, Rojas OL, Rangachari M. Myelin-reactive B cells exacerbate CD4(+) T cell-driven CNS autoimmunity in an IL-23-dependent manner. Nat Commun. 2024;15(1):5404. Epub 20240626. doi: 10.1038/s41467-024-49259-0. PMID: 38926356; PMCID: PMC11208426.

47. Claverie R, Perriguey M, Rico A, Boutiere C, Demortiere S, Durozard P, Hilezian F, Dubrou C, Vely F, Pelletier J, Audoin B, Maarouf A. Efficacy of Rituximab Outlasts B-Cell Repopulation in Multiple Sclerosis: Time to Rethink Dosing? Neurol Neuroimmunol Neuroinflamm. 2023;10(5). Epub 20230821. doi: 10.1212/NXI.0000000000200152. PMID: 37604695; PMCID: PMC10442066.

48. Longbrake EE, Mao-Draayer Y, Cascione M, Zielinski T, Bame E, Brassat D, Chen C, Kapadia S, Mendoza JP, Miller C, Parks B, Xing D, Robertson D. Dimethyl fumarate treatment shifts the immune environment toward an anti-inflammatory cell profile while maintaining protective humoral immunity. Mult Scler. 2021;27(6):883–94. Epub 20200727. doi: 10.1177/1352458520937282. PMID: 32716690; PMCID: PMC8023410.

49. Griffin DO, Holodick NE, Rothstein TL. Human B1 cells in umbilical cord and adult peripheral blood express the novel phenotype CD20+ CD27+ CD43+ CD70. J Exp Med. 2011;208(1):67–80. Epub 20110110. doi: 10.1084/jem.20101499. PMID: 21220451; PMCID: PMC3023138.

50. Yoshitomi H. Peripheral helper T cells, mavericks of peripheral immune responses. Int Immunol. 2024;36(1):9–16. doi: 10.1093/intimm/dxad041. PMID: 37788648; PMCID: PMC10823579.

51. Vermersch P, Granziera C, Mao-Draayer Y, Cutter G, Kalbus O, Staikov I, Dufek M, Saubadu S, Bejuit R, Truffinet P, Djukic B, Wallstroem E, Giovannoni G, Frexalimab Phase 2 Trial G. Inhibition of CD40L with Frexalimab in Multiple Sclerosis. N Engl J Med. 2024;390(7):589–600. doi: 10.1056/NEJMoa2309439. PMID: 38354138.

52. Reidy M, Khan M, Mills EA, Wu Q, Garton J, Draayer DE, Zahoor I, Giri S, Axtell RC, Mao-Draayer Y. New Frontiers in Multiple Sclerosis Treatment: From Targeting Costimulatory Molecules to Bispecific Antibodies. Int J Mol Sci. 2025;26(8). Epub 20250419. doi: 10.3390/ijms26083880. PMID: 40332536; PMCID: PMC12028294.

