## Supplemental Figures for "Ocrelizumab Modulates Both B and T Cell Immune Capacities in Multiple Sclerosis"

**General Cell lineages Gating Strategy**

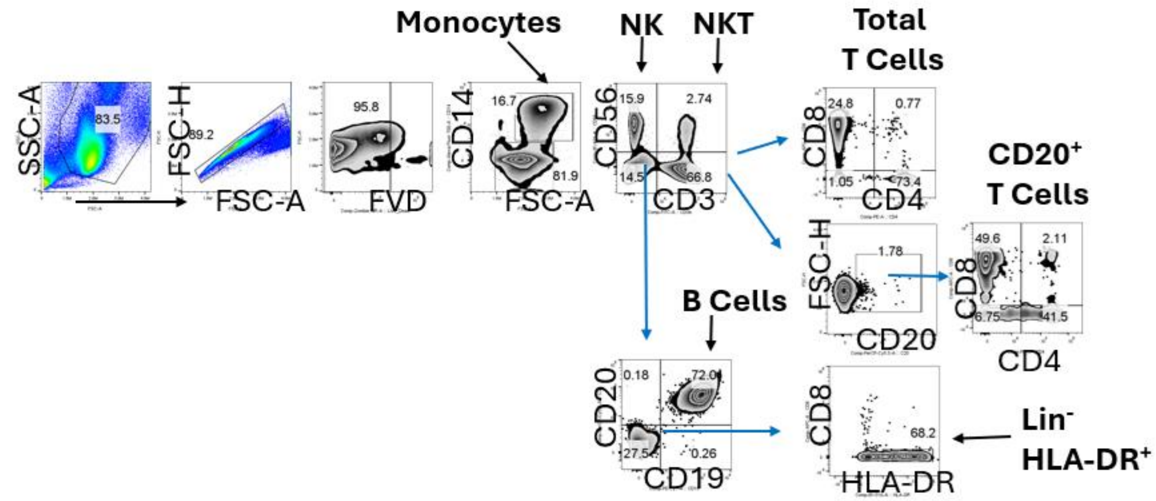


**Gating strategy of Tph**

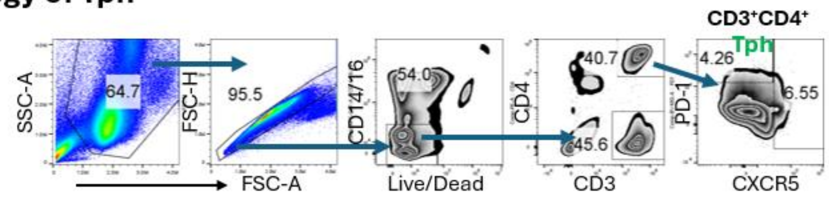


**Supplementary Figure 1. General cell lineage gating strategy.**

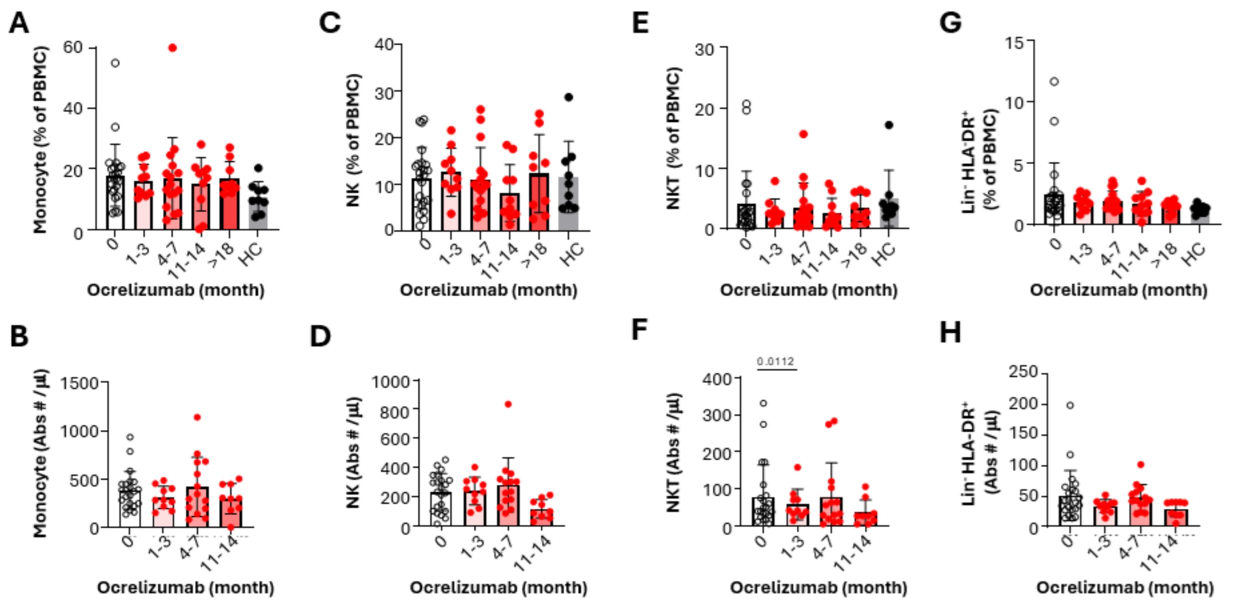


**Supplementary Figure 2. The effects of ocrelizumab on monocytes, NK, NK-T and Lin$^-$HLA-DR$^+$ cells** Effect of frequencies (**A, C, E, G**) and absolute numbers (**B, D, F, H**) of monocytes (FSC-A$^{hi}$, CD14$^+$, **A-B**), NK ( CD14$^-$CD3$^-$CD56$^+$, **C-D),** NKT (CD14$^-$CD3$^+$CD56$^+$, **E-F**) and Lin$^-$(CD14$^-$CD3$^-$CD19$^-$CD8$^-$CD56$^-$)HLA-DR$^+$ cells (**G-H**) were analyzed in PBMC derived from MS patients at baseline (0 month) 1–3 months, 4–7 months, 11–14 months, and beyond 18 months post-treatment. Mixed-effect analysis followed by Dunnet's test for multiple comparisons was used to measure statistical significance between pre-treated and Ocrelizumab-treated MS groups (n=23), while ordinary one-way ANOVA with Dunnet's test for multiple comparisons was used to measure statistical significance between MS pretreatment group with HC (n=9) with $p<0.05$ considered significant.

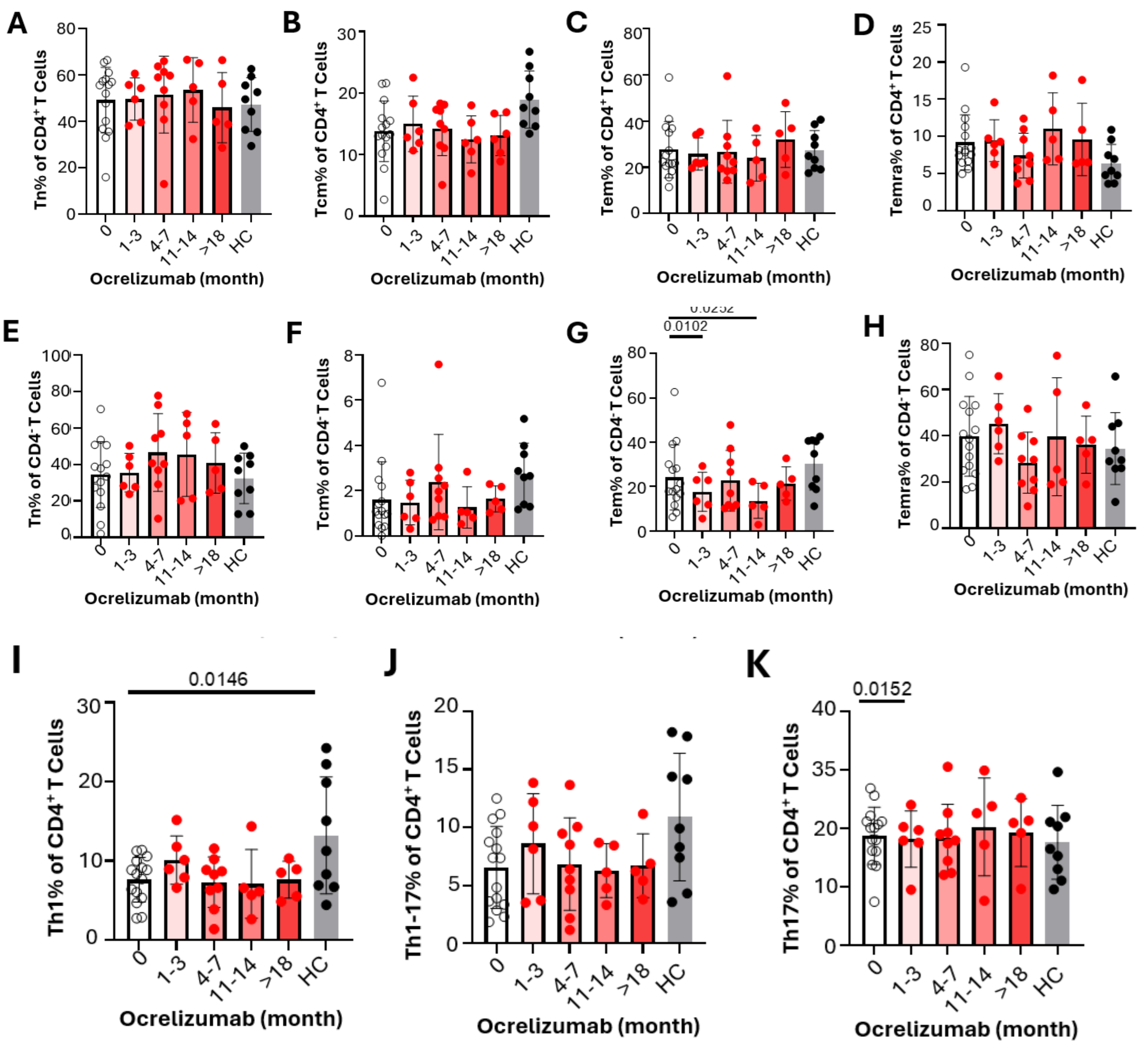


**Supplementary Figure 3. The effect of ocrelizumab on memory and helper T cell subsets. A-D:** memory subsets are shown as the frequencies of $CD4^+$ T cells including $CD45RO^-$,$CCR7^+$ Naïve T cells (Tn, **A**), CD45RO+CCR7+ Central Memory T cells (Tcm, **B**), CD45RO+CCR7- Effector Memory T cells (Tem, **C**) and CD45RO-CCR7- CD45RA+Effector Cells (Temra, **D**). **E-H:** memory subsets are shown as the frequencies of $CD4^-$ T cells including $CD45RO^-CCR7^+$ Naïve T cells (Tn, **E**), CD45RO+CCR7+ Central Memory T cells (Tcm, **F**), $CD45RO^+CCR7^-$ Effector Memory T cells (Tem, **G**) and $CD45RO^-CCR7^-$ $CD45RA^+$Effector Cells (Temra, **H**). **I-K**: T cell helper subsets are shown as frequencies among $CD4^+$ T cells including Th1 ($CXCR3^+$ $CCR6^-CD161^-$, **I**), Th1-17 ($CXCR3^+CCR6^+$ and/or $CD161^+$, **J**) and Th17 ($CXCR3^-CCR6^+$ &/or $CD161^+$, **K**)

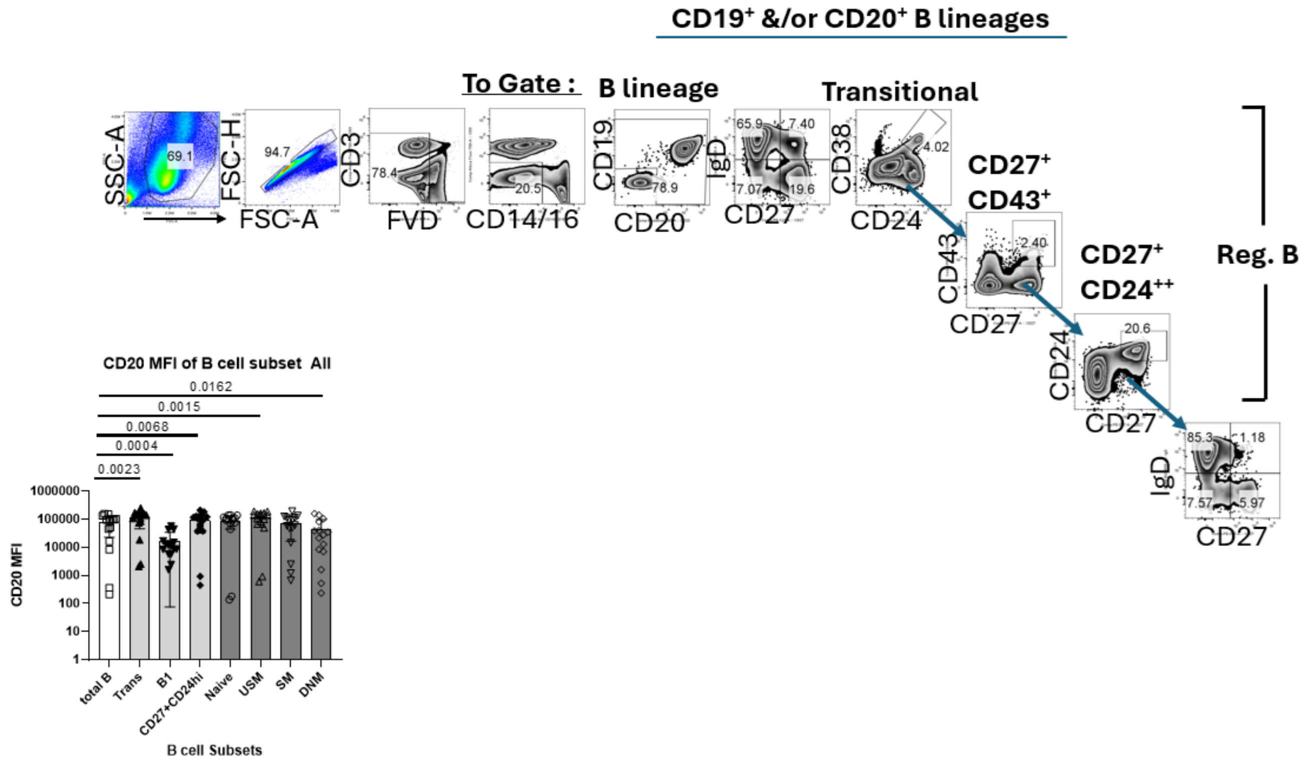


**Supplemental Figure 4. B Cell subset gating strategy and CD20 expression on B cell subsets.** (**A**) B cell subsets analysis gating strategy. (**B**) CD20 expression on subsets of B cells were measured by CD20 mean fluorescence intensity on Total B cells (CD19+ &/or CD20+), Transitional B cells ($CD24^{++}CD38^{++}$), B1 B cells ( $CD27^{+}CD43^{+}$), $CD27^{+}CD24^{hi}$ B cells, as well as rest of Naïve B cells ($CD27^{-l}gD^{+}$), unswitched memory B cells ($CD27^{+}IgD^{+}$), switched memory B cells ($CD27^{+}IgD^{-}$) and double negative B cells ($CD27^{-}IgD^{-}$).

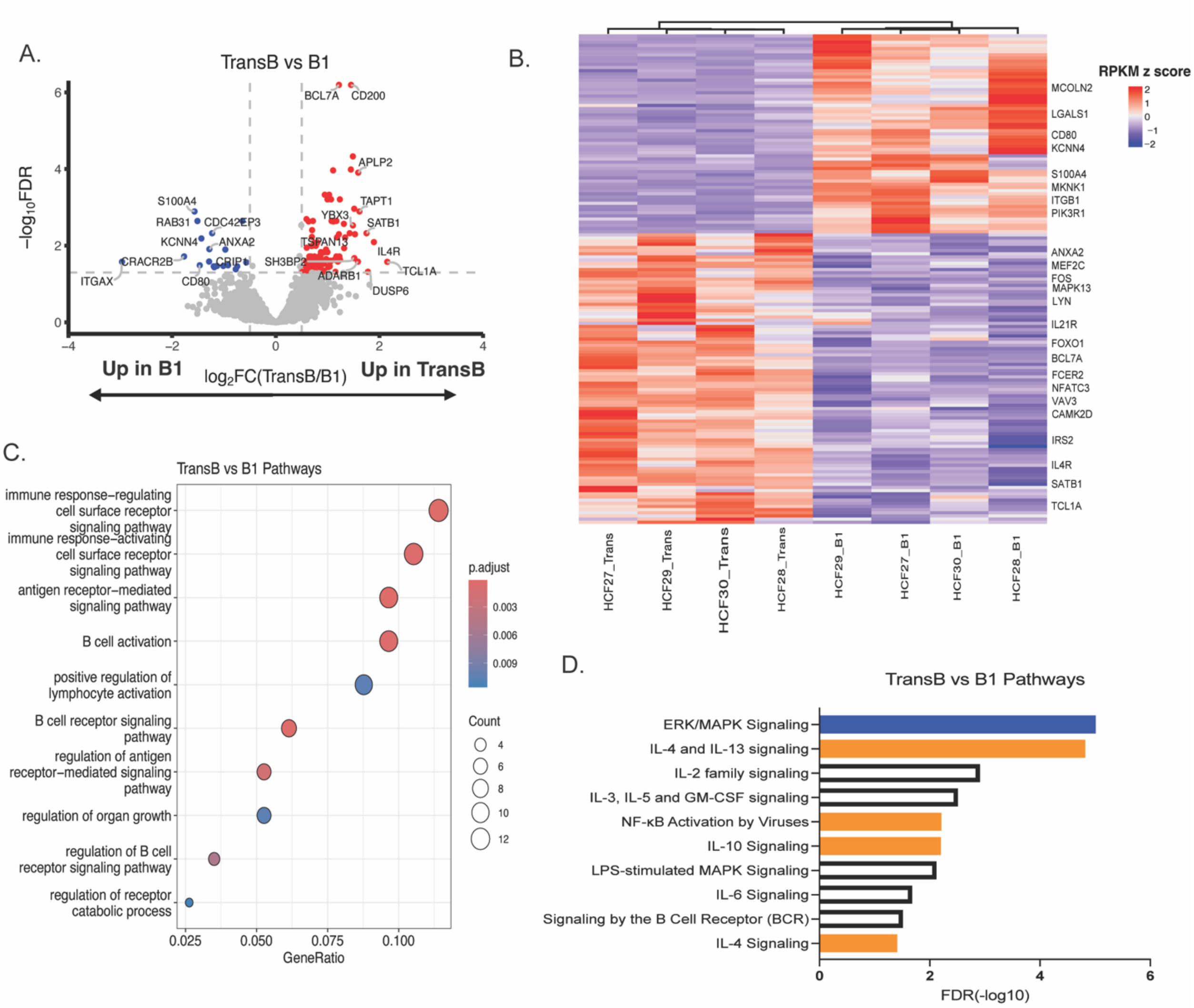


**Supplemental Figure 5. RNA-seq analysis of transitional B vs B1 cells. (A)** Volcano plot of differentially expressed genes (FDR < 0.05, log2FC $\geq \pm 0.5$) between transitional B (n = 4) and B1 (n = 4) cells with the top 10 most significantly up- and down-regulated DEGs shown. Each point denotes a gene expressed in the RNA-seq data. **(B)** Heatmaps of the 131 DEGs identified by differential expression analysis in transitional vs B1 cells. Red color denotes increased expression, and blue color denotes decreased expression. **(C)** Gene Ontology (GO) enrichment analysis of DEGs. The dot plot displays enriched biological process terms. The gene ratio indicates the proportion of DEGs within each GO term. The size of the dot is proportional to the number of genes in the term, and the color gradient represents the adjusted p-value. **(D)** Significant pathways obtained from Ingenuity Pathway Analysis (IPA) of the 169 DEGs in N vs B1. Orange depicts predicted activation; blue indicates predicted inhibition.

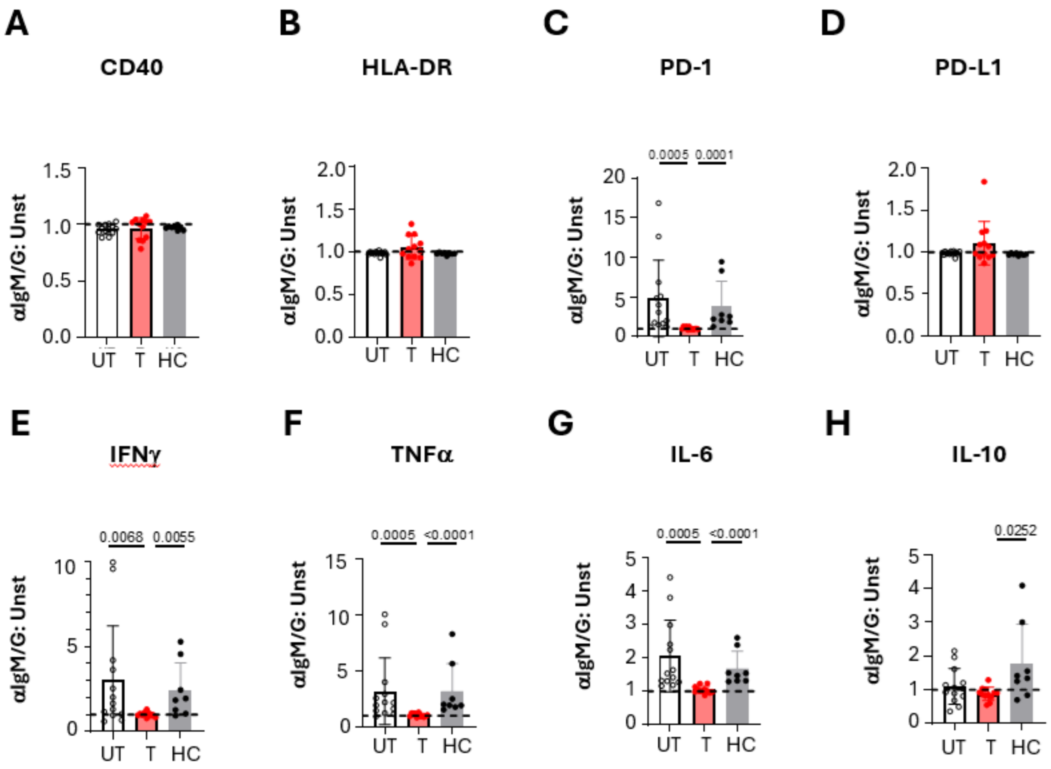


**Supplemental Figure 6**: **Overall effect of ocrelizumab on the functional status of residual B cells**. PBMCs from pre- and post-Ocrelizumab treatment along with heathy controls were incubated for 48hrs with or without anti-IgM/IgG (Fab')2. The ratio of surface molecules (**A-D**) and cytokines (**E-F**) under anti-IgM/G stimulation (aIgM/G) to those under no stimulation (Unst) were analyzed. To explore the overall effect of Ocrelizumab, aggregates of different time points of MS patients post-ocrelizumab treatment (T) were compared with MS before treatment (UT) and healthy control (HC).
